# Competition and cooperation: The plasticity of bacteria interactions across environments

**DOI:** 10.1101/2024.07.03.601864

**Authors:** Josephine Solowiej-Wedderburn, Jennifer T. Pentz, Ludvig Lizana, Björn Schröder, Peter Lind, Eric Libby

**Affiliations:** Department of Mathematics and Mathematical Statistics, Umeå University, Umeå, Sweden; Integrated Science Lab, Umeå University, Umeå, Sweden; Los Alamos National Laboratory, Los Alamos, USA; Department of Physics, Umeå University, Umeå, Sweden; Department of Molecular Biology, Umeå University, Umeå, Sweden; Laboratory for Molecular Infection Medicine Sweden (MIMS), Umeå University, Umeå, Sweden; Umeå Center for Microbial Research (UCMR), Umeå University, Umeå, Sweden

**Keywords:** competition, cooperation, environment, metabolism, microbial ecology

## Abstract

Bacteria live in diverse communities, forming complex networks of interacting species. A central question in bacterial ecology is why some species engage in cooperative interactions, whereas others compete. But this question often neglects the role of the environment. Here, we use genome-scale metabolic networks from two different open-access collections (AGORA and CarveMe) to assess pairwise interactions of different microbes in varying environmental conditions (provision of different environmental compounds). By scanning thousands of environments for 10,000 pairs of bacteria from each collection, we found that most pairs were able to both compete and cooperate depending on the availability of environmental resources. This approach allowed us to determine commonalities between environments that could facilitate the potential for cooperation or competition between a pair of species. Namely, cooperative interactions, especially obligate, were most common in less diverse environments. Further, as compounds were removed from the environment, we found interactions tended to degrade towards obligacy. However, we also found that on average at least one compound could be removed from an environment to switch the interaction from competition to facultative cooperation or vice versa. Together our approach indicates a high degree of plasticity in microbial interactions to the availability of environmental resources.

## Introduction

A core question in ecology is the nature of the interaction between two organisms. For microorganisms such as bacteria this question can be challenging since organisms often interact indirectly by modifying their chemical environment. If two organisms consume the same resources then there can be competition which in simplified systems can drive one species extinct, e.g. competitive exclusion (1). Yet when two organisms secrete resources needed by the other there can be cooperation which can lead to more intimate interactions, e.g. syntrophic networks (2, 3). Within these two extremes there is a large range of interactions and predicting which one actually occurs has important consequences for microbial eco-evolution and our ability to design microbial communities that perform useful functions (4–8). Despite the significant interest in this topic there are still many fundamental questions that remain.

One fundamental question is whether a random pair of microbes is more likely to compete or cooperate. Recent research has increasingly pointed toward competition as the predominant interaction (9). However, investigations into both natural and synthetic microbial communities have also revealed an abundance of cooperative interactions (3, 10, 11). These differences may be explained by the diverse habitats these microbes are sampled from. Different environments may promote distinct interaction patterns. For example, high-resource host-associated environments tend to favour cooperation among species with smaller genomes and high interdependence, whereas free-living habitats tend to favour competition among species with large-genomes and overlapping nutritional requirements (12). Studies also suggest that the types of interactions can be highly dependent on the environment and that even identical pairs of microbes can exhibit vastly different types of interactions (13, 14). For instance, changing the availability of just two resources has been shown to lead to a wide range of different interactions between pairs of auxotrophic microbes that cross-feed by sharing essential amino acids (15). Taken together, these findings suggest that the environment plays a pivotal role in shaping our null expectations regarding microbial interactions.

A prominent example of this phenomenon is the community dynamics of gut microbiota. Within this controlled environment, many species interact and the host can shift interactions by changing the environmental composition. Different studies have suggested that competition (16–18) or cooperation (12, 19, 20) may dominate interactions within the gut. However, a recent study investigating resource competition between gut bacteria found that their model predictions could be improved if cross-feeding was also included, implying a complex interaction landscape with both resource-overlap and cross-feeding potential between species (21). Furthermore, interactions can shift when exposed to different nutrients, as shown recently for communities cultivated on different carbon sources (22). Generally, a variety of dietary fibers maintains a complex and diverse microbial community (23). This is of particular importance when considering the interaction between the gut microbiota and mucus barrier of the host, as fiber fermentation promotes beneficial function of the mucosal barrier (24). Conversely, insufficient fiber consumption can lead to the decline of fiber-restricted specialists in place of generalists, which can switch to the mucus barrier of the host as an alternative energy source (25). Consequently, even in a protective and host-regulated environment such as the gut, substrate availability significantly affects microbial interactions and cross-feeding (11).

Despite the number and complexity of possible interactions between microbes, community assembly experiments have found highly reproducible community structures (26–28). Understanding the role of microbial interactions in driving community assembly is of particular interest and may allow us to make quantitative predictions of how communities should assemble in a given environment. One focus is characterizing the mechanism(s) of cooperative and competitive interactions between members of a community and how these may affect community composition. Cooperative interactions often arise via metabolic cross-feeding which has been shown to emerge between metabolically dissimilar species (29, 30) and can contribute to establishing diverse communities (31, 32). Competitive interactions can occur between metabolically similar species due to competition for available resources and metabolic niche overlap. This has been shown to drive predictable community assembly and structure in both well-controlled laboratory environments (26, 33) as well as more complex environments such as within the gut of the nematode *Caenorhabditis elegans* (18) and on the surface of an *Arabidopsis thaliana* leaf (34). While these studies have shed light on the patterns governing community assembly, it is unclear the extent to which they might be observed outside of controlled lab settings.

We can address the general nature of interactions by adapting tools used to study the microbiome and community assembly. Since a common focus in both has been on the role of metabolic interactions, genome-scale metabolic models provide a useful tool (35, 36). These models contain a set of chemical reactions inferred from genome annotations and can be used to predict growth rates given a set of environmental conditions (usually in terms of supplied chemical compounds). In recent years, metabolic models have been curated for thousands of species with a wide range of applications from the development of personalized medicine (37) to understanding the rarity of prokaryotic endosymbioses (38). Such models have also been used to identify features of different metabolisms that lead to competition or cooperation between species (12, 39–41). In the context of the human gut, these models have been used to explore the effects of different diets on interactions between the microbiota (20, 42) and, more broadly, identify environments which promote e.g. obligate interactions between bacteria (43). By harnessing the power of these metabolic models and the huge databases now available, we can provide null hypotheses for these interactions.

Here, we use metabolic models from two different sources (AGORA (20) and CarveMe (44)) to assess the interactions between thousands of random pairs of bacteria across thousands of different environments. This computational approach allowed us to interrogate the role the environment plays in shaping competition and cooperation on a scale that would be difficult to evaluate empirically. In particular, we varied the number of compounds in the environment to determine the effect of environment diversity on competition and cooperation. By systematically removing compounds from the environment we measured the robustness of interactions to environmental fluctuations and how interactions change as environments degrade. Ultimately, we find that interactions between bacteria show a high degree of plasticity depending on the environment, with a systematic trend towards obligate interactions in less diverse environments.

## Results

We assessed how the environment can shape ecological interactions by using metabolic models from two of the largest open-access collections of genome-scale metabolic networks available: AGORA (20) and CarveMe (44). The AGORA collection includes models for 818 strains of bacteria from the human gut, while CarveMe has models for 5,587 strains of bacteria from disparate places, using genomes from NCBI RefSeq (release 84, (45)). Since the species in AGORA are from the same environment, in contrast to those in CarveMe, we can compare the interactions between bacteria that commonly co-occur relative to those that have may never interacted. In both collections, metabolic models partition metabolites into intracellular and extracellular compartments. We refer to the availability of extracellular compounds as the ‘environment’. By modifying the presence and abundance of these compounds, we can assess the role of the environment on bacterial growth, as computed by the metabolic models.

In both collections, every model comes with a default environment: a set of compounds and concentrations that ensures the growth of the corresponding organism. To determine the nature of an interaction between a bacterial pair, we compared the growth rates of the organisms together versus alone in a new joint environment. The joint environment was created by combining the default environments of the two bacteria such that both were guaranteed to grow. We drew a distinction between two qualitatively different interactions: competitive and cooperative (see Fig. 1A). Competitive interactions are identified when at least one of the organisms must grow slower when both are present. Cooperative interactions are identified when it is possible for both organisms to grow faster when grown together (see *Methods: Assessing growth of individuals and interactions of pairs*). Other interactions are also possible, including neutral interactions which arise when the growth rate of each organism is unaffected by the presence of the other. We considered 10,000 random pairs from each collection and classified their ecological interaction using their joint default environments. We found that the most common default interaction in both collections is neutral (see Fig. 1B), 49% in AGORA and 59% in CarveMe. Of the remaining interactions, we found that the CarveMe collection has more competitive interactions than AGORA and both collections have few cases of cooperation (2% AGORA and 0% CarveMe).

**Fig. 1.**
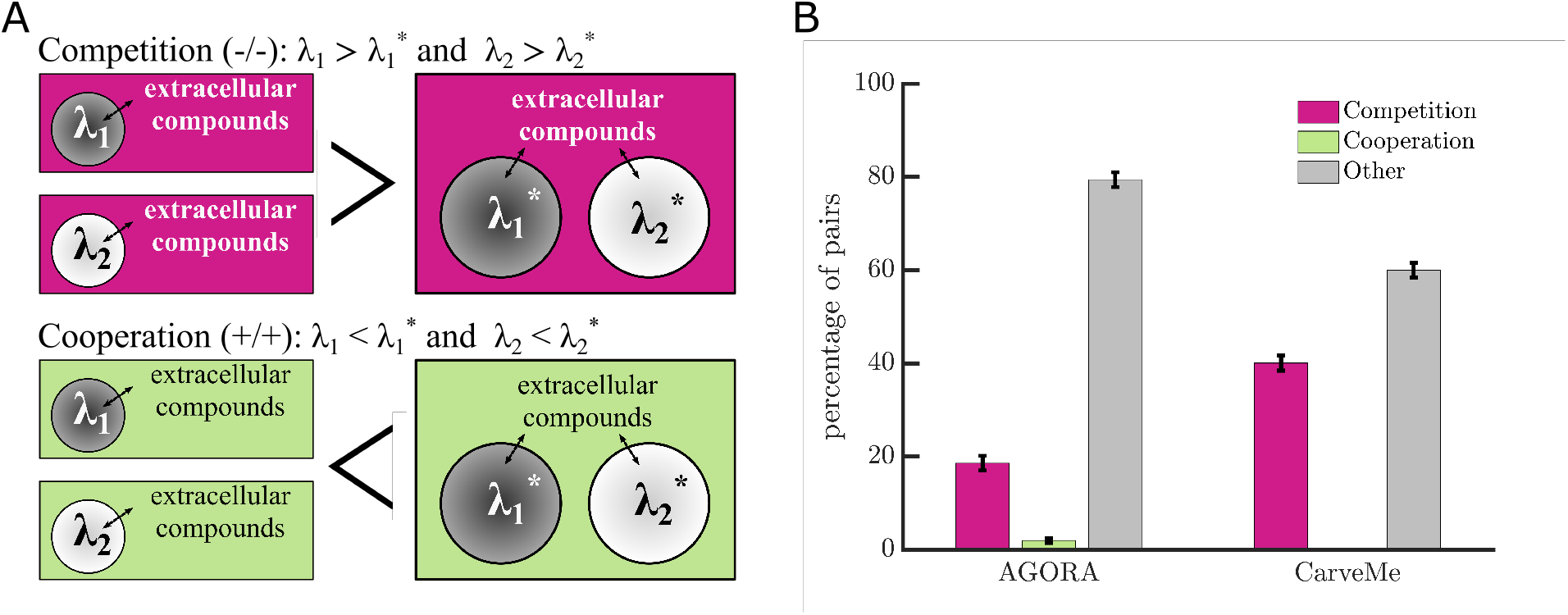
Classifying interactions in default environments. A) A schematic shows interactions between pairs of bacteria (circles) in different environments (rectangles). Competition or cooperation are determined by comparing two computed growth rates for each species in a given environment: 1) the maximal growth rate of each bacteria in isolation (λ_1_ and λ_2_); with 2) the maximal growth rate of each bacteria without harming their partner when grown together (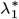 and 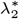, see *Methods: Assessing growth of individuals and interactions of pairs*). We note that with this framework, the same pair of bacteria could have different interactions depending on the environment. B) A bar graph shows the percentage of interactions between pairs of bacteria in their default environments from AGORA and CarveMe. The percentages are the means of 10 samples of 1,000 random pairs. In both collections cooperation is rare in comparison to competition. Over half of the observations were ‘other’ and of these interactions, neutral was most common; the remaining other interactions were commensal (+/=).

To determine whether the interactions present in the default environments were definitive or plastic, we took the same sets of 10,000 pairs of bacteria (from Fig. 1B) and evaluated their interactions in different random environments. We constructed these environments by providing the essential compounds that the bacterial pair must have in any environment in order to grow with no possible substitutes (see *Methods: Essential compounds for pairs*). In addition to the essential compounds, we randomly added a fixed number of environmental compounds that could be utilized by at least one of the bacteria (50 additional compounds for smaller environments; or 100 for larger, see *Methods: Algorithm for generating environments*). For simplicity, we set the availability of all environmental compounds to a fixed but limited amount that is sufficient for growth but can induce competition should both organisms use it (see *Methods: Algorithm for generating environments*). Since the most common interaction in the default environments was neutral, we looked for environments that produced competition or cooperation. For each pair of organisms, we sampled each type of environment until we found 100 viable growth environments. Across all samples, we found at least one cooperative and one competitive environment for 67% (standard deviation 1.2%) of pairs in AGORA (Fig. 2A) and over 87% (standard deviation 0.75%) of pairs in CarveMe (Fig. 2B).

**Fig. 2.**
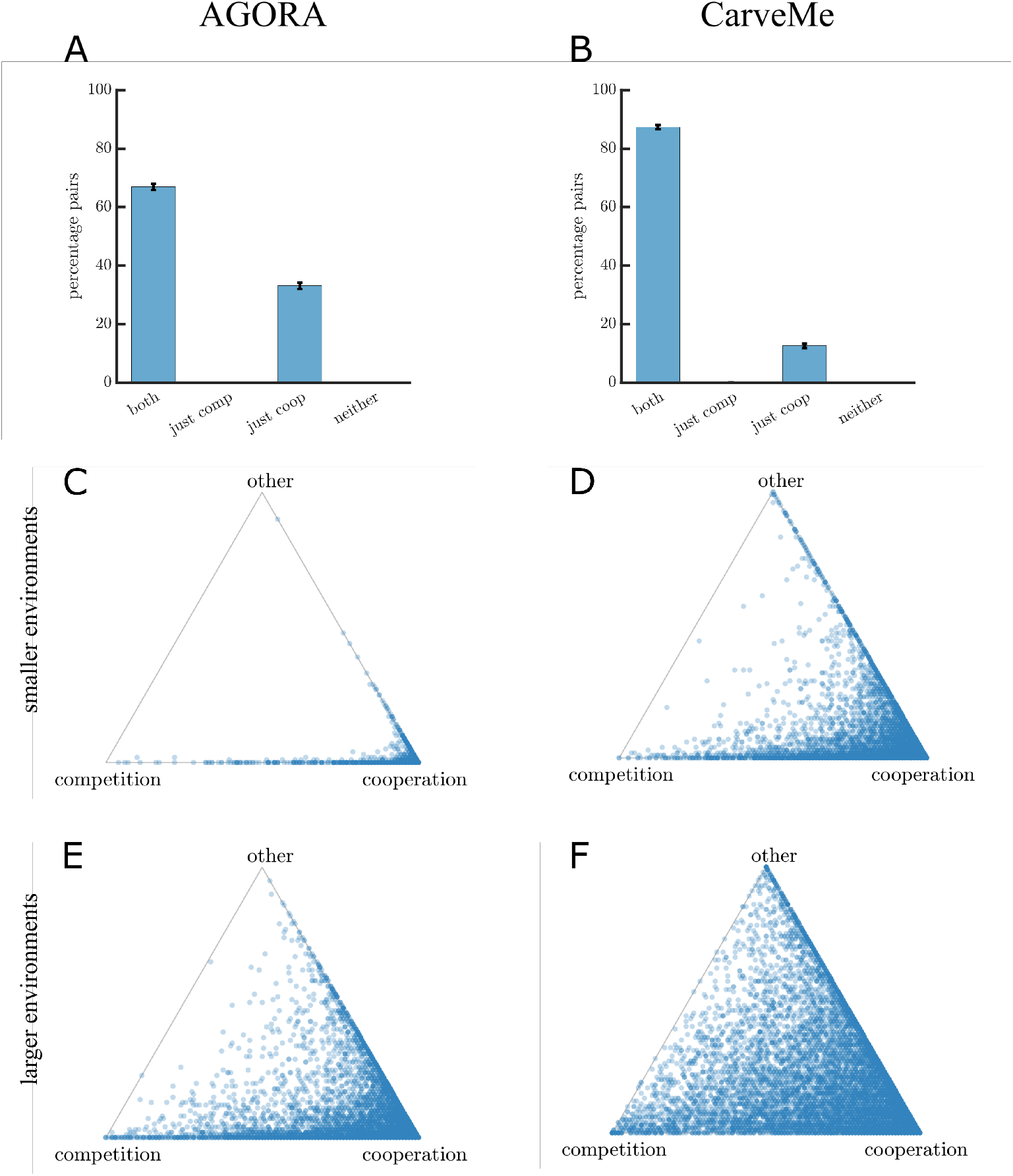
Environment-driven variation in competition and cooperation. A and B) Bar charts show the proportions of bacteria pairs where it was possible to find at least one environment for potential cooperation and/or competition in AGORA (A) and CarveMe (B). The percentages are the means of 10 samples of 1,000 random pairs, as in Fig. 1B. At least one cooperative and one competitive environment were found for 67% (standard deviation 1.2%) pairs in AGORA and 87% pairs (standard deviation 0.8%) in CarveMe; and at least one cooperative environment was found for over 99% pairs across both collections. C–F) Triangle plots show the proportion of different interactions across the 100 viable growth environments identified for pairs of bacteria from (A) and (B). Each point corresponds to a pair of bacteria, with the relative distance to each vertex on the triangle representing the proportion of environments for competitive, cooperative and other interactions identified for that pair. The darkest shading indicates the highest density of points on each plot. Left panel (C and E) shows results for AGORA and right panel (D and F) for CarveMe. Smaller environments (essential compounds plus 50 compounds) are shown in (C) and (D) and larger environments (essential compounds plus 100 compounds) are shown in (E) and (F). We found a more diverse spread of interactions in larger environments.

While the majority of bacterial pairs have the metabolic capacity to both compete and cooperate, the likelihood of finding environments for these interactions differs. Figs. 2C–F show the proportion of environments with competitive, cooperative, or other interactions for different pairs. We observed more examples of cooperative environments than competitive environments across both collections. Consistent with the data from default environments, we found that a higher proportion of environments were competitive in CarveMe in comparison to AGORA. We also found that the relative frequency of observing competitive or cooperative interactions changed as we varied the number of environmental compounds. For example, cooperative interactions were most frequent in the low diversity (small) environments (Figs. 2C and D). Conversely, in high diversity (large) environments, there was a more varied spread of interactions (see Figs. 2E and F). Figs. 2C–F indicate a large degree of variability in the number of competitive or cooperative interactions found between different pairs. Since the underlying organisms represent a wide spectrum of metabolic complexity (e.g., the number and types of reactions present in their metabolic networks), it is possible that some measure of their different metabolic capacities may be predictive of their tendency towards competitive or cooperative interactions. We investigated this possibility by quantifying metabolic complexity in two ways: 1) the number of environmental compounds that a bacteria could use, or 2) the number of metabolic reactions it could perform. For each measure of metabolic complexity, we separated species into three categories based on percentiles: low, medium, and high (see Fig. S4A and B, and S5A and B). In all cases, we rejected the null hypotheses that the number of competitive or cooperative cases was uniform across the different pairings of categories (*p-value* < 0.001, Fig. S4C–F, and S5C–F). We consistently found the fewest cooperative environments between low–low metabolic complexity pairings. While these pairs usually also had the highest number of competitive environments, the results were not always statistically different from pairs of bacteria with other complexity categorizations and, when measuring complexity by number of reactions, the low–low pairs in CarveMe had the fewest competitive environments. We found more cooperative environments between pairs of bacteria where at least one had a high degree of metabolic complexity but the results were not always statistically significant. Overall, these results indicate that there is a difference in the degree of competition or cooperation between bacteria pairings based on their metabolic complexity. However, a more detailed analysis of the metabolic network would be required to be able to predict the variability in competition or cooperation between specific pairs.

If we delve deeper into our data of cooperative and competitive interactions, we observe that there are different types of cooperative interactions. We organized these into three categories: 1) facultative where both microbes can survive alone but have the potential to grow faster together (+/+); 2) oneway obligate where one microbe is reliant on the other and the second microbe can survive alone but benefits from the pairing (× /+); 3) two-way obligate where neither microbe can grow without the other (×/×). Fig. 3 shows that the most common form of cooperation found in our random sampling of different environments was one-way obligate. In ≈ 76% cases (78.3% AGORA and 72.9% CarveMe) the bacteria that was able to survive alone in one-way obligate interactions had a higher metabolic complexity (both when measured by capacity to use environmental compounds or number of reactions). In environments with a high resource diversity (100 additional compounds), the next most frequent type of cooperation was facultative. This differed from environments with a low resource diversity (50 additional compounds) where two-way obligate interactions were the next most frequent type of cooperation observed.

**Fig. 3.**
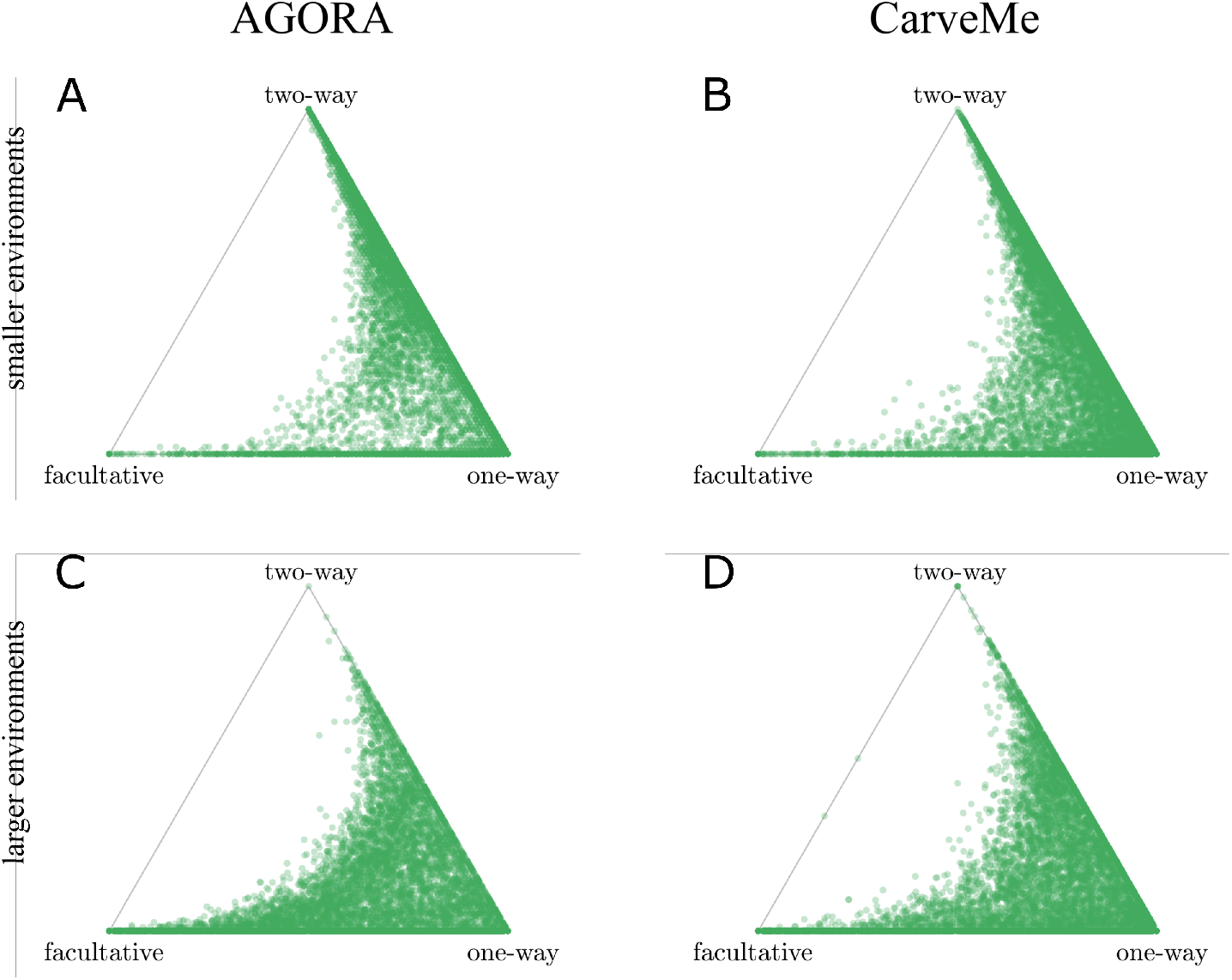
Environment-driven variation in types of cooperation. A–D) Triangle plots show the proportion of different types of cooperative environment: facultative, one-way obligate or two-way obligate, corresponding to pairs of bacteria in Fig. 2C–F. Smaller environments (essential compounds plus 50 compounds) are shown in (A) and (B) and larger environments (essential compounds plus 100 compounds) are shown in (C) and (D). Left panel (A and C) shows results for AGORA and right panel (B and D) for CarveMe. Each point on the plots corresponds to a pair of bacteria, with the relative distance to each vertex on the triangle representing the proportion of environments for each type of cooperative interaction identified for that pair. Where points overlap, the shading is darker, hence the darkest shaded regions have the highest density of points. One-way obligate cooperation was the most common interaction found in both collections and environment sizes. We found more cases of two-way obligate interactions in smaller environments and more cases of facultative cooperation in larger environments.

Given that specific environments facilitate either competitive or cooperative interactions between bacteria, our next step was to investigate how robust these interactions are to perturbations of the environment. We investigated what would happen if we randomly removed a single compound from competitive or cooperative environments for 1,000 random pairs in each collection. In the case of competitive environments, Figs. 4A–D show that the interactions are robust to the loss of single compounds; however, we find the removal of approximately 3–6 of the 100 environmental compounds (means of 6.5 compounds from AGORA and 2.8 from CarveMe) can switch the competitive interaction to cooperative. In the case of cooperative interactions, we focused exclusively on facultative cooperation because obligate interactions cannot lose their obligacy when compounds are removed. We found that, similar to competitive environments, cooperative interactions are robust to the loss of a single compound, though cooperation can switch to competition by removing approximately 1 (means of 1.0 compounds from AGORA and 1.4 from CarveMe) of the compounds (Figs. 4A–D). There are additionally an average of 2-3 compounds (means of 3.5 compounds from AGORA and 2.1 from CarveMe) that cause a transition from facultative to obligate cooperation when removed.

**Fig. 4.**
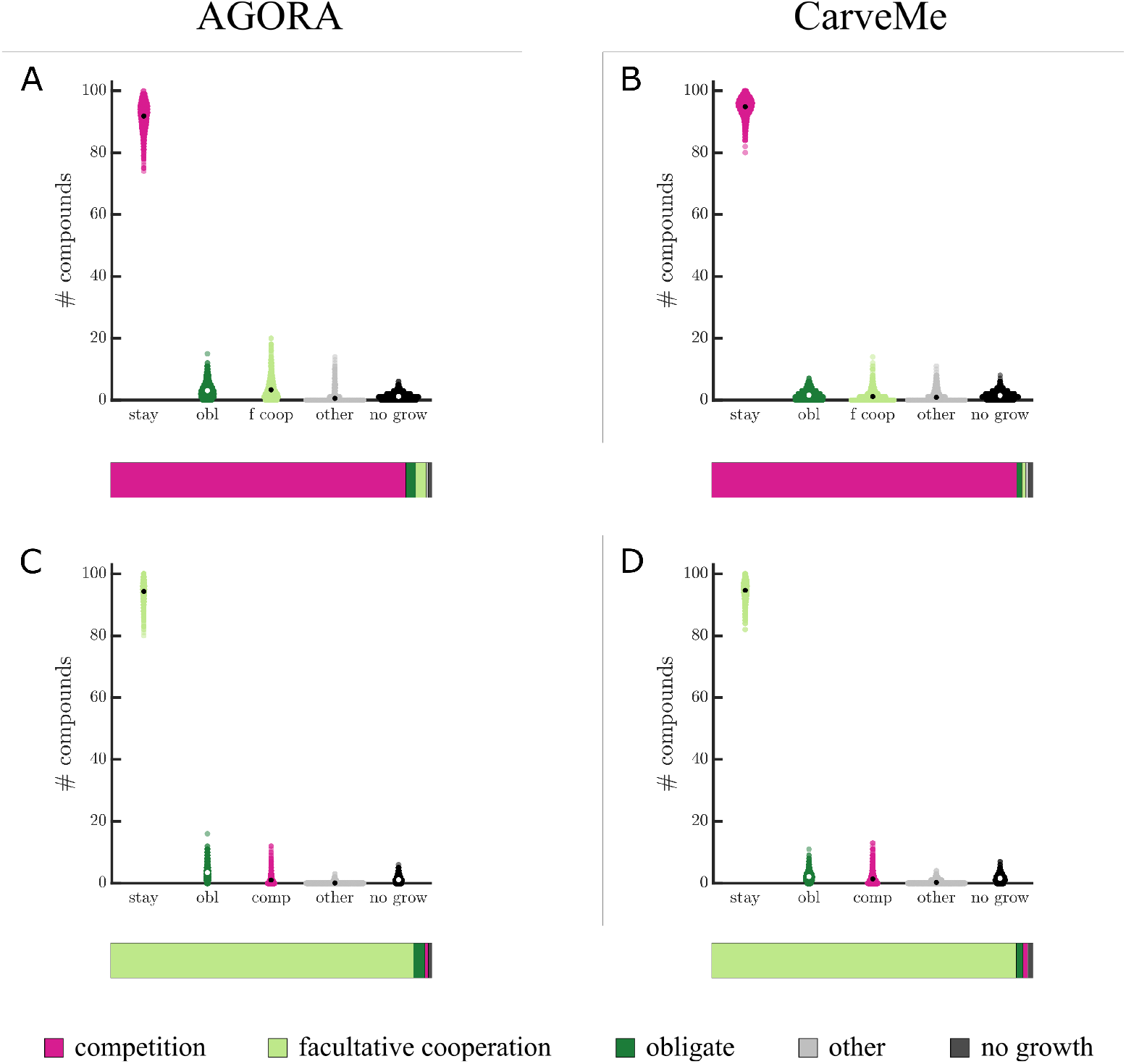
Stability of interactions to the loss of a single compound. A and B) Plots show the frequency that an individual compound could be removed from an initially competitive environment and it would remain competitive, or switch to a different interaction type. Specifically, 1,000 pairs of bacteria in AGORA (A) or CarveMe (B) are considered; for each pair we select one competitive environment with 100 additional compounds and determine the effect when each is individually removed from the starting environment (see *Methods: Single removal of compounds*). Black and white spots indicate the means of each interaction change; summarised in the coloured bars below the plots. In the majority of cases, the interaction would remain competitive if one compound was removed from the environment, however, on average there is at least one compound which would result in a flip from competition to cooperation if removed. E–H) Show the same results starting in environments with facultative cooperation. Qualitatively these appear to mirror the results when starting in competitive environments, for both collections. In Fig. S6 we show that for 70-83% pairs there are 1–6 compounds which can cause a switch to obligate interactions; while for 41–60% pairs there are 1–6 compounds which can cause an interaction switch between competition and facultative cooperation if individually removed.

The previous analyses demonstrated that the difference between competitive and cooperative interactions often depends on a single compound in the environment. Based on this observation, we sought to identify whether any compounds were hallmarks of competitive or cooperative environments. We compared the frequency of compounds in the various environments and found that all compounds appeared in both competitive and cooperative environments (see Fig. S7A and B). We did find that the majority of compounds (> 95%) appeared in a higher proportion of competitive or cooperative environments than by independent random sampling (*p-value*< 0.001); however, the bias was typically small, i.e. the median bias from a proportional representation in competitive and cooperative environments was only 12.1% in AGORA and 8.0% in CarveMe (see Fig. S7C and D).

If we shift our focus to those compounds responsible for switches between competitive and cooperative interactions, we found 45–58% of environmental compounds were responsible for all the interaction switches between competitive and cooperative environments in Figs. 4A–D (45% in AGORA and 58% in CarveMe; results shown in Figs. S8 and S9). Here, the majority of compounds were responsible for fewer than 1% of switches, however, the most frequently observed compounds to cause a switch were in 5–12% of the respective interaction switches, which is statistically significant. Supplemental tables S2 and S3 list the compounds which most commonly caused switches between interactions when removed from the environment. This includes key compounds in energy metabolism: terminal electron acceptors for aerobic (oxygen) and anaerobic (nitrate, nitrite, fumarate) respiration. There is an overlap in compounds that would cause a switch from competitive to cooperative interactions if removed with those that would cause a switch from cooperative to competitive interactions. For example, removing oxygen from the environment commonly caused interaction switches from both competition to cooperation and cooperation to competition between pairs in AGORA. Most of the different categories of compounds identified as relevant for cross-feeding in the gut (11) are also represented here. In particular, several amino acids were found to cause all types of transitions for both databases. Other categories were less universal, for example, B-vitamins (nicotinate (conjugate base of niacin, B3) and riboflavin (B2)) were only common for switches to obligate interactions in AGORA, while switches to obligacy in CarveMe were commonly caused by removal of metal ions (Fe2+, Fe3+ and Cu2+).

We further explored the plasticity of interactions by investigating what happens when environments deteriorate. We did this by systematically removing compounds in competitive and facultative cooperative environments and evaluating whether this altered the type of interaction (see schematic in Fig. 5A and *Methods: Degradation of environments*). We first reduced environments to only those compounds that were being ‘used’ by at least one species, and retaining those known to be essential (see *Methods: Essential compounds for pairs*). Figs. 5B–G show example simulations of pairings of *Prevotella copri* with three other bacteria commonly occurring in the human gut (see Figs. S10–S15 for other pairings between common bacteria in gut, aquatic or soil communities). Each figure shows a set of random sequences of compounds being removed (each column represents a different order of removal), beginning in a particular choice of either a competitive (Figs. 5B–D) or cooperative (Figs. 5E–G) environment. These different pairings highlight some common trends while illustrating the variation caused by the order of compound removal. For example, in the majority of cases that began with a competitive interaction, there was a transition to cooperative interactions. For the choice of initial competitive environment between *Prevotella copri* and *Clostridium perfringens* (Fig. 5C) the switches to cooperation occurred after very few compounds were removed, while in the other pairings more compounds were removed before the switches occurred. Whether starting in a competitive or cooperative environment, the simulations between *Prevotella copri* and *Bacteroides vulgatus* (Figs. 5B and E) show a high frequency of transitions between competition and cooperation, often returning back to the initial interaction, but in the other pairings we typically only observed one or two interaction switches. Finally, we note that the last state before the pairs could no longer grow was often (82%) an obligate interaction.

**Fig. 5.**
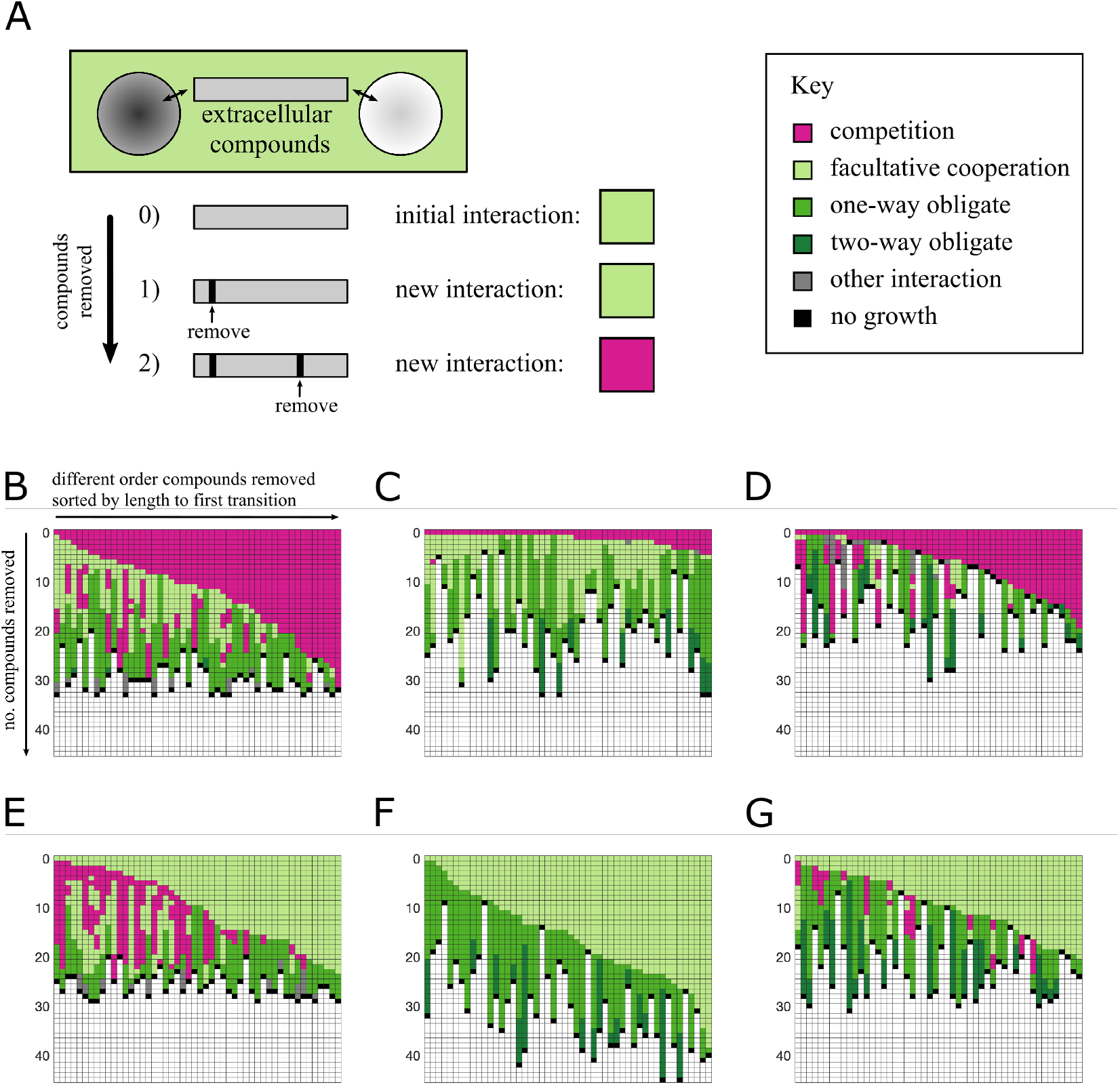
Degrading competitive and cooperative environments for common gut bacteria. A) A schematic shows how compounds are sequentially removed from an environment for facultative cooperation between a pair of bacteria and the resultant change in interaction. Colours correspond to different interactions: competition in pink; facultative cooperation in light green; one-way obligate interactions in green; two-way obligate interactions in dark green; neutral interactions in blue; no growth in black and other interactions in grey ((= / =), (+/ =), or (× / =)). B–G) Heatmaps show how the interactions between three pairs of common gut bacteria change as compounds are removed from the environment (models from AGORA). Going down the columns subsequent compounds are removed from the environment. Each column shows the interaction shifts as different orderings of compounds are removed and columns have been sorted by the number of compounds removed until the first change in interaction. In the top row (B–D) pairs begin in an environment for competition and in the bottom row (E–G) for facultative cooperation. Bacteria pairs *Prevotella copri DSM 18205* and *Bacteroides vulgatus ATCC 8482* (B and E), *Prevotella copri DSM 18205* and *Clostridium perfringens ATCC 13124* (C and F), and *Prevotella copri DSM 18205* and *Ruminococcus gnavus AGR2154* (D and G) show the variety of interaction changes that are commonly observed when environments are reduced. *Prevotella copri* appears to switch more frequently between competition and cooperation when paired with *Bacteroides vulgatus* than *Clostridium perfringens* or *Ruminococcus gnavus*. The most common final interaction before at least one bacteria cannot grow is obligate.

Inspired by these case studies, we considered a larger set of pairs of organisms to see whether the observed trends are consistent. We performed this search for 300 random pairs of bacteria from each collection and starting in either competitive or facultative cooperative environments. Across all these simulations we found that it was common for bacteria to switch to another interaction type before they stopped growing. In AGORA, the competitive environments changed interaction with the removal of fewer compounds than from a cooperative environment (Fig. 6A); while in CarveMe on average, the same number of compounds could be removed from both competitive and cooperative environments before the interaction changed (Fig. 6B). In Figs. 6C–F we summarise the transitions between interactions for 50 different sequences of compounds removed across the different pairings, until at least one of them was no longer able to grow. This shows that, as we found in the case studies, the most common interaction before no growth was obligate. Regardless of the initial interaction, as compounds were removed the most common transition path moved to an obligate interaction and ultimately no growth (left-most path in Figs. 6C–F). The next most common path was to switch between competition and facultative cooperation once before transitioning to an obligate interaction and then ceasing growth. Interestingly, we also observed some sequences with frequent switches, e.g. ≥ 5 transitions between competition and facultative cooperation and vice versa, highlighting the complex relationship between environment and ecology.

**Fig. 6.**
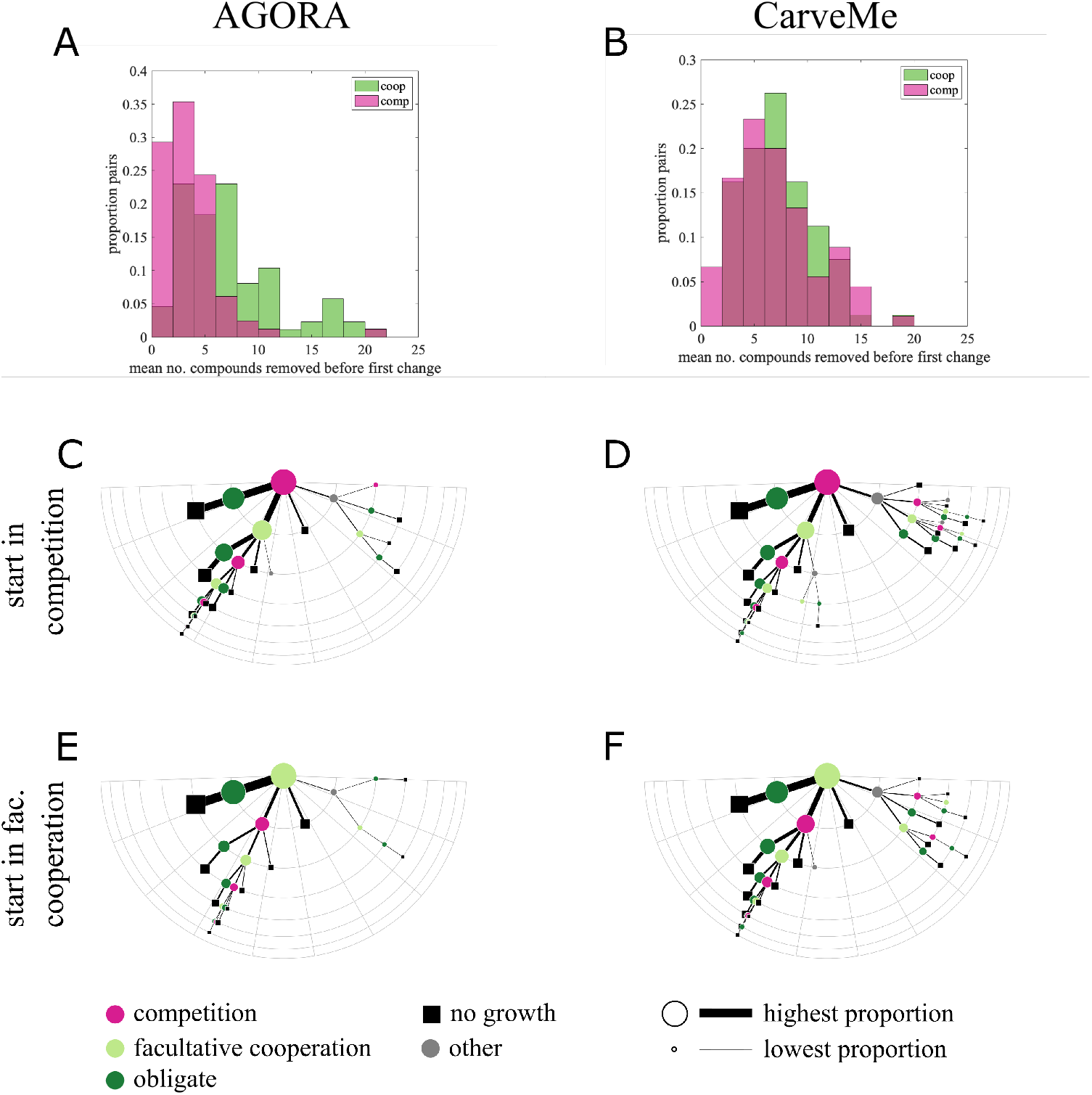
Plasticity of interactions to the loss of environmental compounds. A and B) Histograms show the number of compounds that can be removed before the first change in interaction type for 50 different sequences of compound removal for 300 pairs of bacteria, starting in either competitive or facultative cooperative environments. On average, competitive interactions in AGORA switch with the removal of fewer compounds than cooperative interactions (3.6 *vs*. 7; (A)), while competitive and cooperative interactions both switch after an average removal of 7 compounds in CarveMe (B). C–F) Diagrams summarise the change in interactions between 300 pairs of bacteria as compounds are removed from the environment. The top row summarises interaction paths starting in competition ((C) AGORA and (D) CarveMe), and the bottom starting in facultative cooperation ((E) AGORA and (F) CarveMe). The semi-circular grid indicates the number of interaction switches. Nodes and line thickness are proportional to the square-rooted proportion of interactions taking that path; we only include paths that happened for > 0.1% simulations (see Fig. S16 for full results). Additionally paths are sorted such that the most common branch at each node is furthest to the left. The most common path whether starting in competition or facultative cooperation is to obligate interactions and then no growth.

## Discussion

A fundamental question in ecology is the nature of the interaction between two species. For bacteria these interactions are typically placed on a spectrum between competition and cooperation, with conflicting studies reporting which is more prevalent. While there is an appreciation of some plasticity in specific interactions based on the availability of environmental resources (15), it remains unclear to what extent this applies. Here, we assess the extent to which the environment can modulate ecological interactions by using a large-scale computational approach. By sampling thousands of pairs of bacteria from two different collections of metabolic models, we find that the majority of pairs can exhibit competitive and cooperative interactions, depending on the resources available in the environment (Fig. 2A, B). Despite this variability, we consistently find that cooperation is more common in environments with lower resource diversity. Thus even when pairs initially compete, as resources are consumed and removed pairs switch towards obligate forms of cooperation. These results highlight the dynamic nature of ecological interactions in response to environmental change.

A key result from our analyses is the high degree of plasticity in bacteria interactions depending on the environmental context. Previous studies have found differences in the relative frequencies of cooperative and competitive interactions across different environments (10, 16, 18, 46–48). Since the communities in these studies differ across environmental contexts it is difficult to disentangle whether it is the environment or specific pairs of species that are driving the changes in interactions. One recent study (14) focusing on a specific pair of bacteria, *Agrobacterium tumefaciens* and *Comamonas testosteroni*, has shown that such plasticity is possible as the pair shift between competition and cooperation depending on the concentration of linoleic acid. Through our large-scale metabolic approach we extend this finding and observe that plastic interactions are a common feature of bacteria pairs and should be the null expectation. This plasticity likely stems from the inherent complexity of metabolism in which the average metabolic network can import/export hundreds of different compounds from the environment. This complexity creates diverse opportunities for both competition and cooperation depending on the particular set of compounds available in the environment.

In spite of this complexity, we found some common trends in which environments select for certain types of interactions. Based on the competitive-exclusion principle, one might expect that resource-poor environments will favor competition over the limited availability of compounds (1). At the opposite extreme, resource-rich environments may have a more diverse selection of compounds available for cross-feeding interactions to arise (32). However, our results were more nuanced. In contrast to our initial expectation, we found resource-poor environments were more likely to have cooperative interactions than competition (Figs. 2C and D). Moreover, the cases of competition that we found were more likely to occur in resource-rich environments (Figs. 2C–F). If we consider resource-rich environments, we found cooperation more often than competition but we also found more cases of interactions that were neither competition nor cooperation. Broadly our results align with the stress gradient hypothesis which states that microbes tend to cooperate with each other under more ‘stressful’ conditions but compete more when the stress is reduced (15, 49). While the stress could manifest in different forms, e.g. toxicity (50) or low resource availability (50, 51), in our system they manifest as a low diversity of resources, similar to (22).

Further supporting the stress gradient hypothesis, when compounds were removed from environments it ultimately led to obligate interactions regardless of whether the initial interactions were competitive or cooperative (Fig. 6). This could make sense as poor environments may lack compounds essential for some bacteria that are commonly produced via cross-feeding, such as amino acids (31) or B-vitamins (52, 53). Yet, interestingly, in our random screens of environments obligate interactions were the most common interaction type found with over 99% of bacteria pairs having an obligate interaction in at least one environment. A previous metabolic modeling study also found frequent obligate interactions using a tailored algorithm on a set of 7 species (43). Our analyses generalize this result and demonstrate that even an unbiased random search of environments can readily identify environments where mutual growth is only possible through obligate interactions. The relative ubiquity of obligate interactions has implications for the evolution of biological complexity, e.g. multicellularity or sociality, where obligacy is considered a promising initial state (2, 54, 55).

In our analyses, we found it was difficult to predict whether a particular environment would lead to competition or cooperation. However, the difference between these environments often came down to a single compound. Notably, a significant proportion of all transitions between cooperation and competition occurred through the removal of amino acids, emphasizing their role as modulators of microbial interactions (56, 57). The availability of terminal electron acceptors (e.g., oxygen, nitrate, and nitrite) also played a major role in causing switches, reflecting bacterial diversity in the ability to use anaerobic respiration and fermentation products. We found the removal of other key cross-feeding intermediate metabolites like lacate, succinate, or formate, or short chain fatty acids were less frequent causes of interaction switches, suggesting these might be less universal. Carbon sources, whose role on community composition is often investigated (22, 32), are surprisingly rare among the top compounds initiating interaction switches (Tables S2 and S3; with the exception of benzoate, which has also been highlighted by e.g. (58)). Interestingly, in metabolic models from the AGORA collection, we found many switches to obligacy were due to B-vitamin removal, while in models from the CarveMe collection, Fe2+ and Fe3+ ions were major triggers, reiterating their importance in microbial interactions (52, 53, 59). This highlights that understanding the impact of individual compounds on microbial interactions is essential for regulating community function, as the composition of growth media plays a key role in shaping interactions.

Throughout this paper, we have used metabolic models to assess the effects of environmental resources on the ability of pairs of bacteria to compete or cooperate. While metabolic models have been used in a variety of other contexts (60–66), their development is still an active area of research (37, 67–70). In particular, we note that the models used in this paper do not feature any type of gene regulation. So although we can identify the potential for cooperation or competition what actually happens will depend on a complex array of factors. One way we have tried to mitigate some of the variability from using metabolic approaches was to perform our analyses using two different model collections: AGORA which contains bacteria from the human gut and CarveMe which contains a more diverse collection of bacteria representing metabolisms across all (as sequenced in NCBI RefSeq release 84, (45)) bacterial life. Our main findings concerning the plasticity of bacterial interactions were consistent across the two collections. One instance where the results did systematically differ between collections is in the higher proportion of cooperative interactions observed in AGORA pairings compared with CarveMe. The difference in levels of competition between AGORA and CarveMe pairings is congruent with a recent study that found more cooperative potential between host-associated communities (such as the gut) than in free-living communities (such as in soil) (12, 71). As metabolic network analyses continue to improve, we expect that our ability to draw insights and make predictions for specific pairings will be refined, though we expect our broad conclusions, that were consistent across both collections, are unlikely to change.

Our results suggest a highly dynamic landscape of microbial interactions influenced by the availability of compounds in the environment. This finding has significant implications for our understanding of the eco-evolutionary dynamics of microbial communities. Traditionally, modeling approaches have relied on fixed interactions between pairs of species, such as those described by competitive Lotka-Volterra equations with constant interaction terms. However, our results indicate that a more complex framework, one that accounts for resource fluctuations and their effects on species interactions, may better predict community dynamics and even reveal new types of behaviors (8, 72). The highly plastic nature of these interactions also raises questions about how bacteria navigate them. Specifically, what strategies or heuristics have evolved to help bacteria capitalize on cooperative opportunities? Addressing these questions could illuminate how bacteria regulate their metabolisms to influence their ecological interactions. While our investigations into pairwise interactions provide significant insights into collective behavior, understanding the full impact of the environment on a community requires examining a broader network of interactions. Exploring the constraints and cooperative potential as microbial communities grow presents a fascinating avenue for further investigation.

## Methods

### Assessing growth of individuals and interactions of pairs

We obtained metabolic models of diverse prokaryotes in .xml format made available in ref. (73) for AGORA (20) and CarveMe (44) collections. We followed the procedure outlined in (38) to curate the metabolic models in a format for MATLAB R2022b. These models include a stoichiometric matrix with each column representing a reaction of the metabolism and rows representing chemical compounds (note, the models have been curated such that each row always corresponds to the same compound with names stored in a separate vector). Compounds are separated into two compartments: the extracellular compartment, and intracellular compartment (in AGORA this is simply the cytoplasm, while CarveMe also includes the periplasm separately). Multiplying the stoichiometric matrix by a vector of fluxes for the rate of flow through each reaction determines a righthand-side vector of net changes in compound concentrations in each compartment. Models additionally include upper and lower bounds for all the reaction fluxes and the righthand-side. It is assumed that there can be no accumulation of products in internal compartments and so the upper and lower bounds on the right-hand-side for chemical concentrations in internal compartments are 0, while non-zero bounds on extracellular compounds indicates that their concentration can be altered by the microbes present (note, there are no source or sink reactions in the stoichiometric matrix). Specifically, negative values of the right-hand-side for compounds in the extracellular compartment correspond to a loss from this compartment indicating they are imported by a microbe; positive values correspond to a gain of compounds in the extracellular compartment indicating that they have been exported by a microbe. Hence, we refer to compounds with negative lower bounds in the extracellular compartment as the ‘environment’.

We use the metabolic models to determine the growth rate of microbes in different environments. To do this, we perform flux balance analysis and solve the associated linear program. For example, the linear program for the growth of an individual microbe *M*_*i*_ in isolation is given by

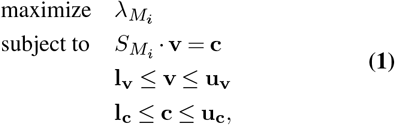

where the objective function, 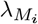, is the flux through the biomass reaction of microbe *M*_*i*_ which is included in the models obtained. In this linear program 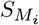 is the stoichiometric matrix of microbe *M*_*i*_; **v** is the vector of reaction fluxes which has lower bounds **l_v_** and upper bounds **u_v_**; the right-hand-side vector **c** is the net change in compound concentrations bounded by **l_c_** and **u_c_**. We solved this and subsequent linear programs by implementing the Gurobi optimization software (74) in MATLAB 2022b.

We determine the ecological interaction of a pair of bacteria in a particular environment by first computing the growth rate of each in isolation. These isolated growth rates serve as reference points against which we compare the growth rates when the bacteria grow together in a shared environment, where they can interact through the import/export of metabolites. The first environment we consider is based on the ‘default’ environment for each organism. In both collections, metabolic models come with an inbuilt default environment—lower and upper bounds on the right-handside vector (**l_c_** and **u_c_**)—designed so that the corresponding microbe can grow in its default environment. By combining the default environments of two microbes we create a new environment that guarantees the growth of the pair and each microbe in isolation. We create this combined environment (as in Fig. 1) by summing the inbuilt values for **l_c_** and **u_c_** for the pair. Implementing these constraints in linear program Eq. (1) determines the growth rates of the individuals, 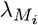 and 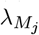.

For a pair, *M*_*i*_ and *M*_*j*_, we construct a combined stoichiometric matrix

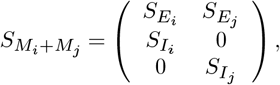

here we have rearranged the rows of 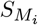 for convenience such that 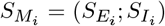 with rows of 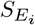 corresponding to compounds in the extracellular compartment and rows of 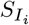 corresponding to the intracellular compartment. We define interactions between organisms by determining their maximal growth rate without harming (reducing the growth rate) of their partner when compared to their individual growth. For example, for pair *M*_*i*_ and *M*_*j*_, we compare the individual growth rate of 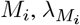, with its growth rate in the shared environment 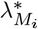, determined by solving the linear program

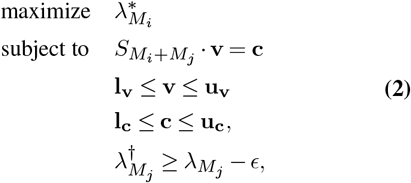

where, 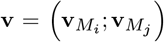 is the combined vector of reaction fluxes each bound the same as for the individuals; 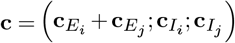 is the right-hand-side vector with a shared extracellular compartment and separate intracellular compartments for *M*_*i*_ and 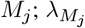 is the individual growth rate determined by solving Eq. (1) for *M*_*j*_ and 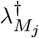 is the growth rate of *M*_*j*_ corresponding to the optimization of *M*_*i*_ when both are growing together. Note, the inequality on the growth rate 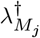 is calculated with a tolerance of ϵ = 0.001. We solve the equivalent linear program for *M*_*j*_ to determine its best possible growth rate, 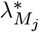, without harming the growth of *M*_*i*_. Table 1 indicates the different possible interactions between microbes *M*_*i*_ and *M*_*j*_. The exploitative interactions (+/-) or (-/+) have been left blank as such interactions are impossible under our optimization scheme: if we have a solution 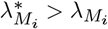 then by construction it must be the case that a 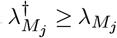 exists which would be a solution for 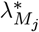.

**Table 1.** Table of potential interaction types for a pair of microbes *M*_*i*_and *M*_*j*_.

| | $\lambda_{M_j}^* < \lambda_{M_j}$ | $\lambda_{M_j}^* = \lambda_{M_j}$ | $\lambda_{M_j}^* > \lambda_{M_j}$ |
| --- | --- | --- | --- |
| $\lambda_{M_i}^* < \lambda_{M_i}$ | (-/-) | (-/=) | |
| $\lambda_{M_i}^* = \lambda_{M_i}$ | (=/-) | (=/=) | (=/+) |
| $\lambda_{M_i}^* > \lambda_{M_i}$ | | (+/=) | (+/+) |

### Essential compounds for pairs

When searching different environments we wanted to minimize the number of environments sampled where the microbes cannot grow together, without otherwise biasing the sampling of environmental compounds. To do so, first we identified a set of environmental compounds that were ‘essential’ to both of the microbes with no possible substitutions. We did this by growing the microbes together in a replete environment where every possible compound in the extracellular compartment was provided (had negative lower bounds). We then individually removed each compound from the replete environment (setting its lower bound to 0) and tested whether both microbes were able to grow together (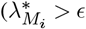 and 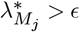, where *ϵ* = 0.1 is a small growth tolerance). If the pair were not able to grow together when a particular compound was removed, this compound was identified as ‘essential’ as no possible substitutions exist to replace it. When constructing environments, we always provided the essential compounds for that pair. However, we note that the set of essential compounds was not necessarily sufficient—other compounds might be necessary for the growth of the microbe pair but potential substitutions exist which need to be provided. See Fig. S1 for a histogram of the number of essential compounds per pair and table S1 for the 10 most common essential compounds across the 10,000 pairs we considered.

### Algorithm for generating environments

We generated different environments by changing the bounds of the right-hand-side vector **c** (**l_c_** and **u_c_**). As positive values of **c** indicate the export of compounds into the environment, we kept the upper bounds on the right-hand-side as the sum of default values for the pair throughout all our analyses. Changing the lower bounds on **c** we were able to explore the effects of the availability of different environmental compounds on microbial interactions.

In this analysis we focused on the presence and absence of compounds in the environment and their effect on the interaction between pairs of microbes. As such, when generating environments, we set **l_c_** = − 1000mmol gDW^− 1^ h^− 1^ for compounds present in the environment; this is the default value for a single microbe and so deemed to be a value sufficient for growth but with the potential to induce competition between the pairs. By contrast, if both microbes included the same compound in their default environments, we would set **l_c_** = − 2000mmol gDW^− 1^ h^− 1^ when considering the interaction in their combined default environment. Compounds not present in the environment had **l_c_** = 0. We additionally repeated the analysis for 10,000 pairs with a reduced concentration of 500mmol gDW^− 1^ h^− 1^ and obtained similar results (see Fig. S3). Of course, further lowering the concentration of compounds, we expect to generate higher degrees of competition between pairs.

Each pair was always provided with its essential compounds (identified by *Methods: Essential compounds for pairs*). In addition, we randomly sampled a fixed number of additional compounds (50 or 100) uniformly from those extracellular compounds that could be used by at least one of the pair. These usable compounds were identified as those which appeared in at least one reaction in the combined stoichiometric matrix. On average, there were 177 usable compounds per pair in AGORA and 206 usable compounds per pair in CarveMe (see Fig. S2), hence we sampled fixed size environments of 50 or 100 additional compounds. When selecting 10,000 random pairings in each collection, we had to discard 317 randomly selected pairs from AGORA and 58 from CarveMe as their combined environment included less than 100 compounds once the essential compounds had been removed.

### Single removal of compounds

To determine the stability of interactions to the loss of a single compound from a given environment for a particular pair of bacteria, we computed the interaction when each of the additional compounds was individually removed from the environment. As essential compounds are essential to that pair in any environment (with no possible substitutes), we did not remove these compounds. We went through each of the additional compounds one-by-one removing them from the complete starting environment (essential compounds plus all 100 additional compounds) and determining the effect of removing it from the environment on the interaction between the pair. This process was repeated for each compound.

### Degradation of environments

To determine the sensitivity of interactions to perturbations of the environment, we first examined whether the interaction between 1,000 different pairs would change upon the removal of any single compound from either competitive or facultative cooperative environments. We then wanted to explore the plasticity of interactions between pairs by removing the additional (substitutable) compounds from both competitive and cooperative starting environments until at least one organism was no longer able to grow. As only some of the available compounds are imported by the pair (i.e. only some of the environmental compounds have negative values for **c**_*E*_ when the linear problem Eq. (2) is solved), we removed any compounds from the initial environment that were not imported by either of the microbes. From this reduced environment, we then sequentially removed compounds until at least one organism was no longer able to grow. We did not remove essential compounds as it was already known that this would terminate the sequence. (Note, that while the other compounds are not essential-without-substitutes, once the environment is reduced such that no substitute remains, at least one of the microbes can no longer grow.) For each pair considered, we tried two different starting environments, one competitive and one facultative cooperative, and 50 different random orderings of compound removal.

## ACKNOWLEDGEMENTS

JSW acknowledges funding from Kempestiftelserna (grant JCK-2129.2). This work has been cleared for public release by the Los Alamos National Laboratory LA-UR-24-26155. This project has been further funded through the Swedish Research Council (Vetenskapsrådet, grant 2021-06602).

## Bibliography

1. Garrett Hardin. The competitive exclusion principle: an idea that took a century to be born has implications in ecology, economics, and genetics. science, 131(3409):1292–1297, 1960.

2. Glen D’Souza, Shraddha Shitut, Daniel Preussger, Ghada Yousif, Silvio Waschina, and Christian Kost. Ecology and evolution of metabolic cross-feeding interactions in bacteria. Natural product reports, 35(5):455–488, 2018.

3. Christian Kost, Kiran Raosaheb Patil, Jonathan Friedman, Sarahi L. Garcia, and Markus Ralser. Metabolic exchanges are ubiquitous in natural microbial communities. Nature Microbiology, pages 1–9, November 2023. ISSN 2058-5276. doi: 10.1038/s41564-023-01511-x. Publisher: Nature Publishing Group.

4. Katharine Z Coyte, Jonas Schluter, and Kevin R Foster. The ecology of the microbiome: networks, competition, and stability. Science, 350(6261):663–666, 2015.

5. Karen De Roy, Massimo Marzorati, Pieter Van den Abbeele, Tom Van de Wiele, and Nico Boon. Synthetic microbial ecosystems: an exciting tool to understand and apply microbial communities. Environmental microbiology, 16(6):1472–1481, 2014.

6. Michael T Mee and Harris H Wang. Engineering ecosystems and synthetic ecologies. Molecular BioSystems, 8(10):2470–2483, 2012.

7. Alvaro Sanchez, Djordje Bajic, Juan Diaz-Colunga, Abigail Skwara, Jean CC Vila, and Seppe Kuehn. The community-function landscape of microbial consortia. Cell Systems, 14(2):122–134, 2023.

8. Feng Liu, Junwen Mao, Wentao Kong, Qiang Hua, Youjun Feng, Rashid Bashir, and Ting Lu. Interaction variability shapes succession of synthetic microbial ecosystems. Nature communications, 11(1):309, 2020.

9. Jacob D. Palmer and Kevin R. Foster. Bacterial species rarely work together. Science, 376 (6593):581–582, May 2022. ISSN 0036-8075, 1095-9203. doi: 10.1126/science.abn5093.

10. Jared Kehe, Anthony Ortiz, Anthony Kulesa, Jeff Gore, Paul C. Blainey, and Jonathan Friedman. Positive interactions are common among culturable bacteria. Science Advances, 7 (45):eabi7159, November 2021. doi: 10.1126/sciadv.abi7159. Publisher: American Association for the Advancement of Science.

11. Elizabeth J. Culp and Andrew L. Goodman. Cross-feeding in the gut microbiome: Ecology and mechanisms. Cell Host & Microbe, 31(4):485–499, April 2023. ISSN 1931-3128. doi: 10.1016/j.chom.2023.03.016. Publisher: Elsevier.

12. Daniel Machado, Oleksandr M. Maistrenko, Sergej Andrejev, Yongkyu Kim, Peer Bork, Kaustubh R. Patil, and Kiran R. Patil. Polarization of microbial communities between competitive and cooperative metabolism. Nature Ecology & Evolution, 5(2):195–203, February 2021. ISSN 2397-334X. doi: 10.1038/s41559-020-01353-4. Number: 2 Publisher: Nature Publishing Group.

13. Yizhu Qiao, Qiwei Huang, Hanyue Guo, Meijie Qi, He Zhang, Qicheng Xu, Qirong Shen, and Ning Ling. Nutrient status changes bacterial interactions in a synthetic community. Applied and Environmental Microbiology, 90(1):e01566–23, December 2023. doi: 10.1128/aem.01566-23. Publisher: American Society for Microbiology.

14. Rita Di Martino, Aurore Picot, and Sara Mitri. Oxidative stress changes interactions between 2 bacterial species from competitive to facilitative. PLOS Biology, 22(2):e3002482, February 2024. ISSN 1545-7885. doi: 10.1371/journal.pbio.3002482. Publisher: Public Library of Science.

15. Tim A. Hoek, Kevin Axelrod, Tommaso Biancalani, Eugene A. Yurtsev, Jinghui Liu, and Jeff Gore. Resource Availability Modulates the Cooperative and Competitive Nature of a Microbial Cross-Feeding Mutualism. PLOS Biology, 14(8):e1002540, August 2016. ISSN 1545-7885. doi: 10.1371/journal.pbio.1002540. Publisher: Public Library of Science.

16. Ophelia S Venturelli, Alex V Carr, Garth Fisher, Ryan H Hsu, Rebecca Lau, Benjamin P Bowen, Susan Hromada, Trent Northen, and Adam P Arkin. Deciphering microbial interactions in synthetic human gut microbiome communities. Molecular Systems Biology, 14(6): e8157, June 2018. ISSN 1744-4292. doi: 10.15252/msb.20178157. Publisher: John Wiley & Sons, Ltd.

17. Karoline Faust, Franziska Bauchinger, Béatrice Laroche, Sophie de Buyl, Leo Lahti, Alex D. Washburne, Didier Gonze, and Stefanie Widder. Signatures of ecological processes in microbial community time series. Microbiome, 6(1):120, June 2018. ISSN 2049-2618. doi: 10.1186/s40168-018-0496-2.

18. Anthony Ortiz, Nicole M. Vega, Christoph Ratzke, and Jeff Gore. Interspecies bacterial competition regulates community assembly in the c. elegans intestine. Proceedings of the National Academy of Science, 15:2131—-2145, 2021. doi: 10.1038/s41396-021-00910-4.

19. Sebastian Germerodt, Katrin Bohl, Anja Lück, Samay Pande, Anja Schröter, Christoph Kaleta, Stefan Schuster, and Christian Kost. Pervasive Selection for Cooperative Cross-Feeding in Bacterial Communities. PLOS Computational Biology, 12(6):e1004986, June 2016. ISSN 1553-7358. doi: 10.1371/journal.pcbi.1004986. Publisher: Public Library of Science.

20. Stefanía Magnúsdóttir, Almut Heinken, Laura Kutt, Dmitry A. Ravcheev, Eugen Bauer, Alberto Noronha, Kacy Greenhalgh, Christian Jäger, Joanna Baginska, Paul Wilmes, Ronan M. T. Fleming, and Ines Thiele. Generation of genome-scale metabolic reconstructions for 773 members of the human gut microbiota. Nature Biotechnology, 35(1):81–89, January 2017. ISSN 1546-1696. doi: 10.1038/nbt.3703. Number: 1 Publisher: Nature Publishing Group.

21. Po-Yi Ho, Taylor H. Nguyen, Juan M. Sanchez, Brian C. DeFelice, and Kerwyn Casey Huang. Resource competition predicts assembly of gut bacterial communities in vitro. Nature Microbiology, 9(4):1036–1048, April 2024. ISSN 2058-5276. doi: 10.1038/s41564-024-01625-w. Publisher: Nature Publishing Group.

22. Erin Ostrem Loss, Jaron Thompson, Pak Lun Kevin Cheung, Yili Qian, and Ophelia S. Venturelli. Carbohydrate complexity limits microbial growth and reduces the sensitivity of human gut communities to perturbations. Nature Ecology & Evolution, 7(1):127–142, January 2023. ISSN 2397-334X. doi: 10.1038/s41559-022-01930-9. Number: 1 Publisher: Nature Publishing Group.

23. Kassem Makki, Edward C. Deehan, Jens Walter, and Fredrik Bäckhed. The Impact of Dietary Fiber on Gut Microbiota in Host Health and Disease. Cell Host & Microbe, 23(6): 705–715, June 2018. ISSN 19313128. doi: 10.1016/j.chom.2018.05.012.

24. Sandra M. Holmberg, Rachel H. Feeney, Vishnu Prasoodanan P. K. Fabiola Puértolas-Balint, Dhirendra K. Singh, Supapit Wongkuna, Lotte Zandbergen, Hans Hauner, Beate Brandl, Anni I. Nieminen, Thomas Skurk, and Bjoern O. Schroeder. The gut commensal Blautia maintains colonic mucus function under low-fiber consumption through secretion of short-chain fatty acids. Nature Communications, 15(1):3502, April 2024. ISSN 2041-1723. doi: 10.1038/s41467-024-47594-w. Publisher: Nature Publishing Group.

25. Mahesh S. Desai, Anna M. Seekatz, Nicole M. Koropatkin, Nobuhiko Kamada, Christina A. Hickey, Mathis Wolter, Nicholas A. Pudlo, Sho Kitamoto, Nicolas Terrapon, Arnaud Muller, Vincent B. Young, Bernard Henrissat, Paul Wilmes, Thaddeus S. Stappenbeck, Gabriel Núñez, and Eric C. Martens. A Dietary Fiber-Deprived Gut Microbiota Degrades the Colonic Mucus Barrier and Enhances Pathogen Susceptibility. Cell, 167(5):1339–1353.e21, November 2016. ISSN 00928674. doi: 10.1016/j.cell.2016.10.043.

26. Joshua E. Goldford, Nanxi Lu, Djordje Bajić, Sylvie Estrela, Mikhail Tikhonov, Alicia Sanchez-Gorostiaga, Daniel Segrè, Pankaj Mehta, and Alvaro Sanchez. Emergent simplicity in microbial community assembly. Science, 361:469–474, 2018. doi: 10.1126/science.aat1168.

27. Sylvie Estrela, Jean C.C. Vila, Nanxi Lu, Djordje Bajić, Maria Rebolleda-Gómez, Chang-Yu Chang, Joshua E. Goldford, Alicia Sanchez-Gorostiaga, and Álvaro Sánchez. Functional attractors in microbial community assembly. Cell Systems, 13:29–42, 2021. doi: 10.1016/j.cels.2021.09.011.

28. He Fu, Mario Uchimiya, Jeff Gore, and Mary Ann Moran. Ecological drivers of bacterial community assembly in synthetic phycospheres. Proceedings of the National Academy of Science, 117:3656–3662, 2020. doi: 10.1073/pnas.1917265117.

29. Hsuan-Chao Chiu, Roie Levy, and Elhanan Borenstein. Emergent Biosynthetic Capacity in Simple Microbial Communities. PLOS Computational Biology, 10(7):e1003695, July 2014. ISSN 1553-7358. doi: 10.1371/journal.pcbi.1003695. Publisher: Public Library of Science.

30. Samir Giri, Leonardo Oña, Silvio Waschina, Shraddha Shitut, Ghada Yousif, Christoph Kaleta, and Christian Kost. Metabolic dissimilarity determines the establishment of crossfeeding interactions in bacteria. Current Biology, 31(24):5547–5557.e6, December 2021. ISSN 0960-9822. doi: 10.1016/j.cub.2021.10.019.

31. Leonardo Oña, Samir Giri, Neele Avermann, Maximilian Kreienbaum, Kai M. Thormann, and Christian Kost. Obligate cross-feeding expands the metabolic niche of bacteria. Nature Ecology & Evolution, 5(9):1224–1232, September 2021. ISSN 2397-334X. doi: 10.1038/s41559-021-01505-0. Number: 9 Publisher: Nature Publishing Group.

32. Martina Dal Bello, Hyunseok Lee, Akshit Goyal, and Jeff Gore. Resource–diversity relationships in bacterial communities reflect the network structure of microbial metabolism. Nature Ecology & Evolution, 5(10):1424–1434, October 2021. ISSN 2397-334X. doi: 10.1038/s41559-021-01535-8. Number: 10 Publisher: Nature Publishing Group.

33. Hyunseok Lee, Blox Bloxham, and Jeff Gore. Resource competition can explain simplicity in microbial community assembly. Proceedings of the National Academy of Science, 120, 2023. doi: 10.1073/pnas.2212113120.

34. Martin Schäfer, Alan R. Pacheco, Rahel Künzler, Miriam Bortfeld-Miller, Christopher M. Field, Evangelia Vayena, Vassily Hatzimanikatis, and Julia A. Vorholt. Metabolic interaction models recapitulate leaf microbiota ecology. Science, 381, 2023. doi: 10.1126/science.adf5121.

35. Eugen Bauer and Ines Thiele. From Network Analysis to Functional Metabolic Modeling of the Human Gut Microbiota. mSystems, 3(3):e00209–17, March 2018. doi: 10.1128/mSystems.00209-17. Publisher: American Society for Microbiology.

36. Lillian R. Dillard, Dawson D. Payne, and Jason A. Papin. Mechanistic models of microbial community metabolism. Molecular Omics, 17(3):365–375, 2021. doi: 10.1039/D0MO00154F. Publisher: Royal Society of Chemistry.

37. Almut Heinken, Johannes Hertel, Geeta Acharya, Dmitry A. Ravcheev, Malgorzata Nyga, Onyedika Emmanuel Okpala, Marcus Hogan, Stefanía Magnúsdóttir, Filippo Martinelli, Bram Nap, German Preciat, Janaka N. Edirisinghe, Christopher S. Henry, Ronan M. T. Fleming, and Ines Thiele. Genome-scale metabolic reconstruction of 7,302 human microorganisms for personalized medicine. Nature Biotechnology, pages 1–12, January 2023. ISSN 1546-1696. doi: 10.1038/s41587-022-01628-0. Publisher: Nature Publishing Group.

38. Eric Libby, Christopher P. Kempes, and Jordan G. Okie. Metabolic compatibility and the rarity of prokaryote endosymbioses. Proceedings of the National Academy of Sciences, 120(17):e2206527120, April 2023. doi: 10.1073/pnas.2206527120. Publisher: Proceedings of the National Academy of Sciences.

39. Shiri Freilich, Raphy Zarecki, Omer Eilam, Ella Shtifman Segal, Christopher S. Henry, Martin Kupiec, Uri Gophna, Roded Sharan, and Eytan Ruppin. Competitive and cooperative metabolic interactions in bacterial communities. Nature Communications, 2(1):589, December 2011. ISSN 2041-1723. doi: 10.1038/ncomms1597. Number: 1 Publisher: Nature Publishing Group.

40. Roie Levy and Elhanan Borenstein. Metabolic modeling of species interaction in the human microbiome elucidates community-level assembly rules. Proceedings of the National Academy of Sciences, 110(31):12804–12809, July 2013. doi: 10.1073/pnas.1300926110. Publisher: Proceedings of the National Academy of Sciences.

41. Aleksej Zelezniak, Sergej Andrejev, Olga Ponomarova, Daniel R. Mende, Peer Bork, and Kiran Raosaheb Patil. Metabolic dependencies drive species co-occurrence in diverse microbial communities. Proceedings of the National Academy of Sciences, 112(20):6449–6454, May 2015. doi: 10.1073/pnas.1421834112. Publisher: Proceedings of the National Academy of Sciences.

42. Almut Heinken and Ines Thiele. Anoxic Conditions Promote Species-Specific Mutualism between Gut Microbes In Silico. Applied and Environmental Microbiology, 81(12):4049–4061, June 2015. doi: 10.1128/AEM.00101-15. Publisher: American Society for Microbiology.

43. Niels Klitgord and Daniel Segrè. Environments that Induce Synthetic Microbial Ecosystems. PLOS Computational Biology, 6(11):e1001002, November 2010. ISSN 1553-7358. doi: 10.1371/journal.pcbi.1001002. Publisher: Public Library of Science.

44. Daniel Machado, Sergej Andrejev, Melanie Tramontano, and Kiran Raosaheb Patil. Fast automated reconstruction of genome-scale metabolic models for microbial species and communities. Nucleic Acids Research, 46(15):7542–7553, September 2018. ISSN 1362-4962. doi: 10.1093/nar/gky537.

45. Kim D. Pruitt, Tatiana Tatusova, and Donna R. Maglott. NCBI reference sequences (Ref-Seq): a curated non-redundant sequence database of genomes, transcripts and proteins. Nucleic Acids Research, 35(suppl_1):D61–D65, January 2007. ISSN 0305-1048. doi: 10.1093/nar/gkl842.

46. Anna S Weiss, Anna G Burrichter, Abilash Chakravarthy Durai Raj, Alexandra von Strempel, Chen Meng, Karin Kleigrewe, Philipp C Münch, Luis Rössler, Claudia Huber, Wolfgang Eisenreich, et al. In vitro interaction network of a synthetic gut bacterial community. The ISME journal, 16(4):1095–1109, 2022.

47. Charlotte I Carlström, Christopher M Field, Miriam Bortfeld-Miller, Barbara Müller, Shinichi Sunagawa, and Julia A Vorholt. Synthetic microbiota reveal priority effects and keystone strains in the arabidopsis phyllosphere. Nature Ecology & Evolution, 3(10):1445–1454, 2019.

48. Kevin R Foster and Thomas Bell. Competition, not cooperation, dominates interactions among culturable microbial species. Current biology, 22(19):1845–1850, 2012.

49. Erik F. Y. Hom and Andrew W. Murray. Plant-fungal ecology. Niche engineering demonstrates a latent capacity for fungal-algal mutualism. Science (New York, N.Y.), 345(6192): 94–98, July 2014. ISSN 1095-9203. doi: 10.1126/science.1253320.

50. Philippe Piccardi, Björn Vessman, and Sara Mitri. Toxicity drives facilitation between 4 bacterial species. Proceedings of the National Academy of Sciences, 116(32):15979–15984, August 2019. doi: 10.1073/pnas.1906172116. Publisher: Proceedings of the National Academy of Sciences.

51. Sarah P. Hammarlund and William R. Harcombe. Refining the stress gradient hypothesis in a microbial community. Proceedings of the National Academy of Sciences, 116(32):15760–15762, August 2019. doi: 10.1073/pnas.1910420116. Publisher: Proceedings of the National Academy of Sciences.

52. Rachel Gregor, Rachel Emoke Szabo, Gabriel Toneatti Vercelli, Matti Gralka, Ryan Reynolds, Evan B Qu, Naomi M Levine, and Otto X Cordero. Widespread b-vitamin auxotrophy in marine particle-associated bacteria. bioRxiv, pages 2023–10, 2023.

53. Stefanía Magnúsdóttir, Dmitry Ravcheev, Valérie de Crécy-Lagard, and Ines Thiele. Systematic genome assessment of b-vitamin biosynthesis suggests co-operation among gut microbes. Frontiers in genetics, 6:129714, 2015.

54. Sylvie Estrela, Benjamin Kerr, and J Jeffrey Morris. Transitions in individuality through symbiosis. Current opinion in microbiology, 31:191–198, 2016.

55. Eric Libby and William C Ratcliff. Lichens and microbial syntrophies offer models for an interdependent route to multicellularity. The Lichenologist, 53(4):283–290, 2021.

56. Svenja Starke, Danielle MM Harris, Johannes Zimmermann, Sven Schuchardt, Mhmd Oumari, Derk Frank, Corinna Bang, Philip Rosenstiel, Stefan Schreiber, Norbert Frey, et al. Amino acid auxotrophies in human gut bacteria are linked to higher microbiome diversity and long-term stability. The ISME Journal, 17(12):2370–2380, 2023.

57. Jason SL Yu, Clara Correia-Melo, Francisco Zorrilla, Lucia Herrera-Dominguez, Mary Y Wu, Johannes Hartl, Kate Campbell, Sonja Blasche, Marco Kreidl, Anna-Sophia Egger, et al. Microbial communities form rich extracellular metabolomes that foster metabolic interactions and promote drug tolerance. Nature microbiology, 7(4):542–555, 2022.

58. Susse Kirkelund Hansen, Paul B Rainey, Janus AJ Haagensen, and Søren Molin. Evolution of species interactions in a biofilm community. Nature, 445(7127):533–536, 2007.

59. Jos Kramer, Özhan Özkaya, and Rolf Kümmerli. Bacterial siderophores in community and host interactions. Nature Reviews Microbiology, 18(3):152–163, 2020.

60. Maxime Durot, Pierre-Yves Bourguignon, and Vincent Schachter. Genome-scale models of bacterial metabolism: reconstruction and applications. FEMS microbiology reviews, 33(1): 164–190, 2008.

61. William R Harcombe, William J Riehl, Ilija Dukovski, Brian R Granger, Alex Betts, Alex H Lang, Gracia Bonilla, Amrita Kar, Nicholas Leiby, Pankaj Mehta, et al. Metabolic resource allocation in individual microbes determines ecosystem interactions and spatial dynamics. Cell reports, 7(4):1104–1115, 2014.

62. Sergey Stolyar, Steve Van Dien, Kristina Linnea Hillesland, Nicolas Pinel, Thomas J Lie, John A Leigh, and David A Stahl. Metabolic modeling of a mutualistic microbial community. Molecular systems biology, 3(1):92, 2007.

63. Eric Libby, Laurent Hébert-Dufresne, Sayed-Rzgar Hosseini, and Andreas Wagner. Syntrophy emerges spontaneously in complex metabolic systems. PLoS computational biology, 15(7):e1007169, 2019.

64. Joshua E Goldford, Hyman Hartman, Temple F Smith, and Daniel Segrè. Remnants of an ancient metabolism without phosphate. Cell, 168(6):1126–1134, 2017.

65. Harrison B Smith, Alexa Drew, John F Malloy, and Sara Imari Walker. Seeding biochemistry on other worlds: Enceladus as a case study. Astrobiology, 21(2):177–190, 2021.

66. João F Matias Rodrigues and Andreas Wagner. Evolutionary plasticity and innovations in complex metabolic reaction networks. PLoS computational biology, 5(12):e1000613, 2009.

67. Changdai Gu, Gi Bae Kim, Won Jun Kim, Hyun Uk Kim, and Sang Yup Lee. Current status and applications of genome-scale metabolic models. Genome biology, 20(1):1–18, 2019.

68. Johan Gustafsson, Mihail Anton, Fariba Roshanzamir, Rebecka Jörnsten, Eduard J Kerkhoven, Jonathan L Robinson, and Jens Nielsen. Generation and analysis of context-specific genome-scale metabolic models derived from single-cell rna-seq data. Proceedings of the National Academy of Sciences, 120(6):e2217868120, 2023.

69. Can Chen, Chen Liao, and Yang-Yu Liu. Teasing out missing reactions in genome-scale metabolic networks through hypergraph learning. Nature Communications, 14(1):2375, 2023.

70. Christian Diener and Sean M Gibbons. More is different: metabolic modeling of diverse microbial communities. Msystems, 8(2):e01270–22, 2023.

71. Seth Rakoff-Nahoum, Kevin R Foster, and Laurie E Comstock. The evolution of cooperation within the gut microbiota. Nature, 533(7602):255–259, 2016.

72. Samuel FM Hart, Hanbing Mi, Robin Green, Li Xie, Jose Mario Bello Pineda, Babak Momeni, and Wenying Shou. Uncovering and resolving challenges of quantitative modeling in a simplified community of interacting cells. PLoS biology, 17(2):e3000135, 2019.

73. Christian Lieven, Moritz E. Beber, Brett G. Olivier, Frank T. Bergmann, Meric Ataman, Parizad Babaei, Jennifer A. Bartell, Lars M. Blank, Siddharth Chauhan, Kevin Correia, Christian Diener, Andreas Dräger, Birgitta E. Ebert, Janaka N. Edirisinghe, José P. Faria, Adam M. Feist, Georgios Fengos, Ronan M. T. Fleming, Beatriz García-Jiménez, Vassily Hatzimanikatis, Wout van Helvoirt, Christopher S. Henry, Henning Hermjakob, Markus J. Herrgård, Ali Kaafarani, Hyun Uk Kim, Zachary King, Steffen Klamt, Edda Klipp, Jasper J. Koehorst, Matthias König, Meiyappan Lakshmanan, Dong-Yup Lee, Sang Yup Lee, Sunjae Lee, Nathan E. Lewis, Filipe Liu, Hongwu Ma, Daniel Machado, Radhakrishnan Mahadevan, Paulo Maia, Adil Mardinoglu, Gregory L. Medlock, Jonathan M. Monk, Jens Nielsen, Lars Keld Nielsen, Juan Nogales, Intawat Nookaew, Bernhard O. Palsson, Jason A. Papin, Kiran R. Patil, Mark Poolman, Nathan D. Price, Osbaldo Resendis-Antonio, Anne Richelle, Isabel Rocha, Benjamín J. Sánchez, Peter J. Schaap, Rahuman S. Malik Sheriff, Saeed Shoaie, Nikolaus Sonnenschein, Bas Teusink, Paulo Vilaça, Jon Olav Vik, Judith A. H. Wodke, Joana C. Xavier, Qianqian Yuan, Maksim Zakhartsev, and Cheng Zhang. MEMOTE for standardized genome-scale metabolic model testing. Nature Biotechnology, 38(3):272–276, March 2020. ISSN 1546-1696. doi: 10.1038/s41587-020-0446-y. Publisher: Nature Publishing Group.

74. Gurobi Optimization, LLC. Gurobi Optimizer Reference Manual, 2021.

