## Supplementary Information for "Competition and cooperation: The plasticity of bacteria interactions across environments"

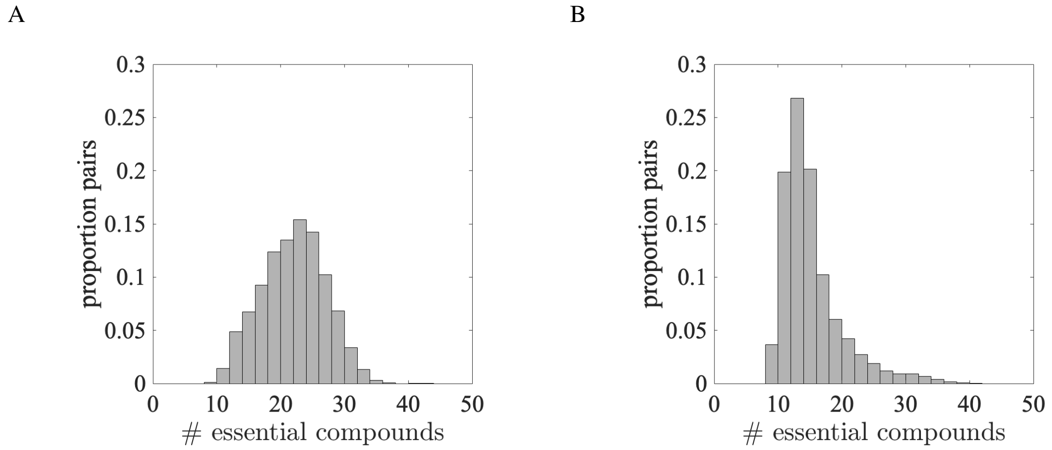


**Fig. S1. Essential compounds per pair.** Histograms show the number of essential compounds for 10,000 pairs in AGORA (A) and CarveMe (B) collections, as used in the main paper.

**Table S1. Common essential compounds.** The 10 most common essential compounds for pairs of bacteria in AGORA (a) and CarveMe (b).

**(a)**

| Compound | % pairs |
|---|---|
| zinc | 100% |
| manganese | 100% |
| magnesium | 100% |
| potassium | 100% |
| copper (2+) | 100% |
| cobalt (2+) | 100% |
| chloride | 100% |
| calcium (2+) | 100% |
| sulfate | 97.2% |
| thiamin | 89.4% |

**(b)**

| Compound | % pairs |
|---|---|
| zinc | 100% |
| magnesium | 100% |
| potassium | 100% |
| cobalt (2+) | 100% |
| chloride | 100% |
| calcium (2+) | 100% |
| manganese | 99.9% |
| oxygen | 89.6% |
| sulfate | 82.0% |
| citrate | 52.1% |

DRAFT

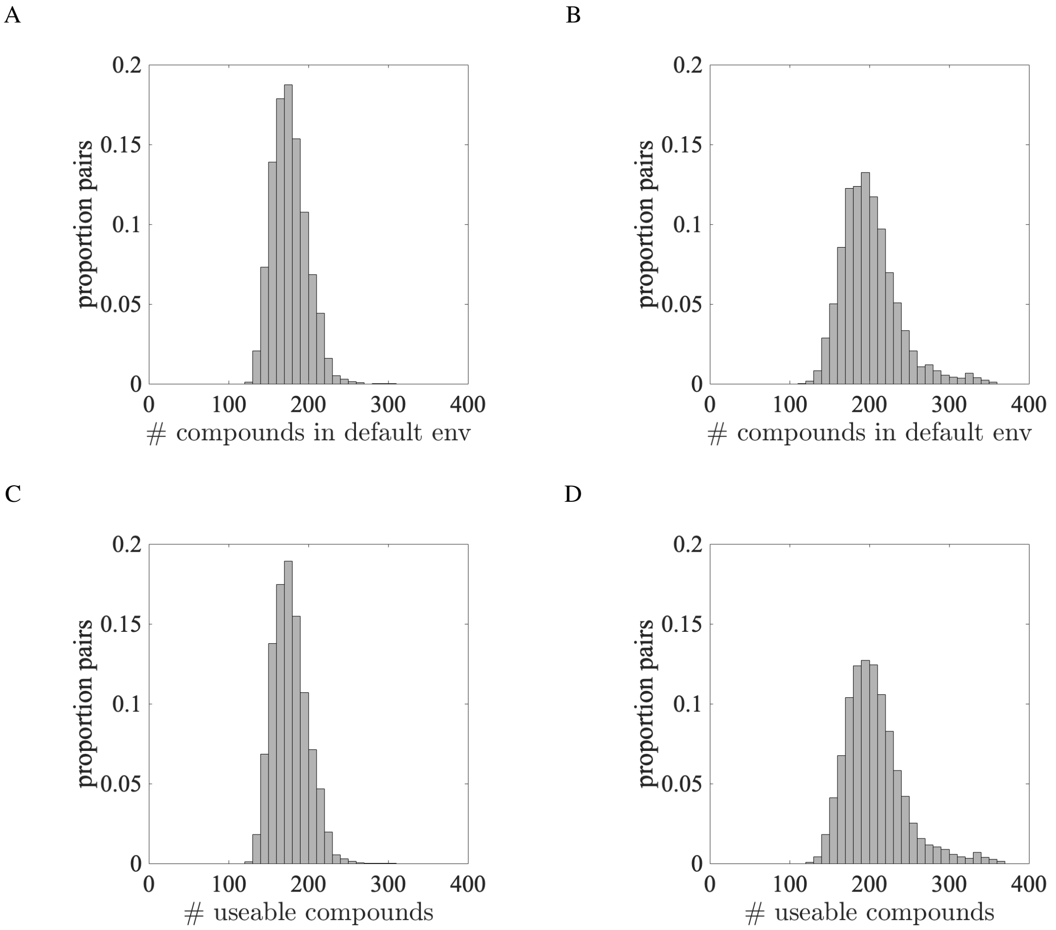


**Fig. S2. Potential size of environments per pair.** A and B) Histograms show the number of environmental compounds in combined default environments for 10,000 pairs in AGORA (A) and CarveMe (B) collections, as used in the main paper Fig. 1B. C and D) Histograms show the total number of environmental compounds that could be used by the same 10,000 pairs in AGORA (C) and CarveMe (D) collections. This is the number of environmental compounds that at least one of the bacteria contain within a metabolic reaction. Default environments contain almost all useable environmental compounds hence the histograms (A) and (B) look similar to (C) and (D).

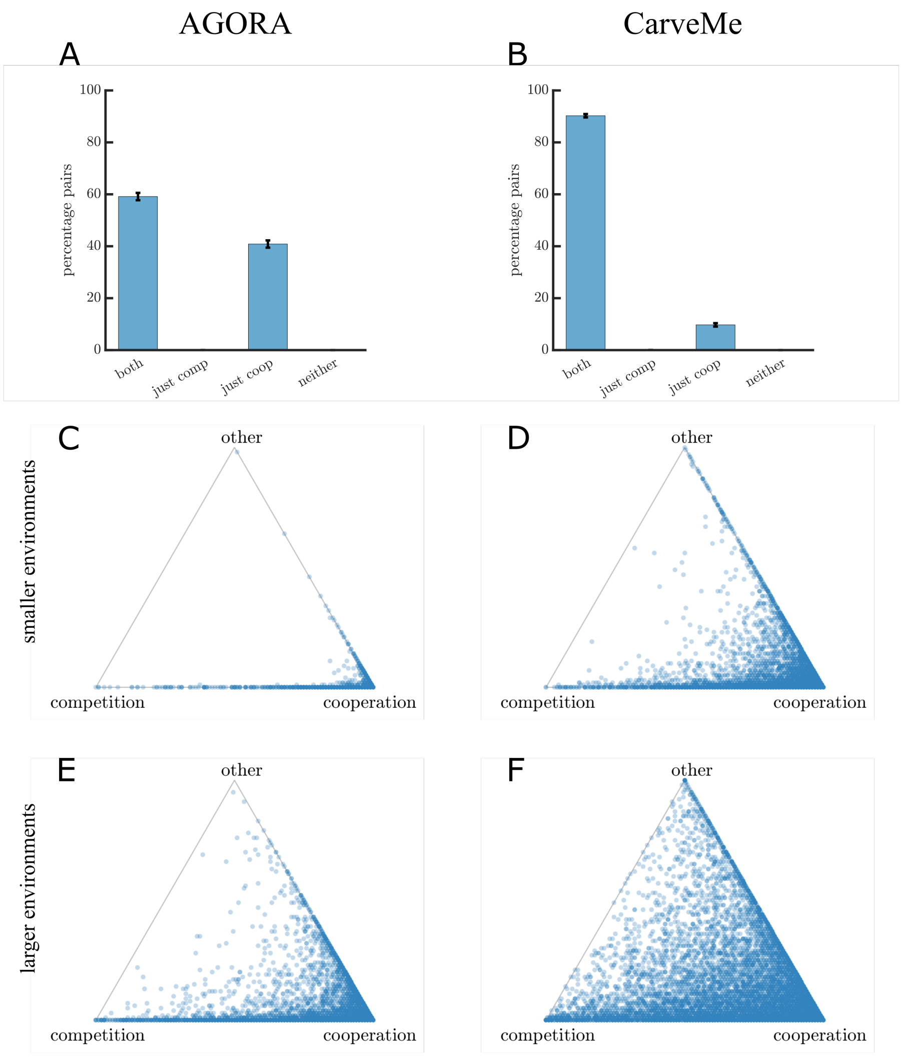


**Fig. S3. Environment-driven variation in competition and cooperation with available concentration flux -500mmol gDW$^{-1}$ h$^{-1}$** Figure shows the same analysis as Fig. 2 in the main paper but with a reduced value for the lower bound of the concentration flux (right-hand-side of the flux balance analysis) of environmentally available compounds; where we set the right-hand-side lower bound to -1000mmol gDW$^{-1}$ h$^{-1}$ in the main paper, we here use -500mmol gDW$^{-1}$ h$^{-1}$. A and B) Bar charts show the proportions of bacteria pairs where it was possible to find at least one environment for potential cooperation and/or competition in AGORA (A) and CarveMe (B). We now find at least one competitive and one cooperative environment for 59% (standard deviation 1.6%) pairs in AGORA and 90% pairs (standard deviation 0.9%) in CarveMe. C–F) Triangle plots show the proportion of different interactions across the 100 viable growth environments identified for pairs of bacteria from (A) and (B). Each point corresponds to a pair of bacteria, with the relative distance to each vertex on the triangle representing the proportion of environments for competitive, cooperative and other interactions identified for that pair. The darkest shading indicates the highest density of points on each plot. Left panel (C and E) shows results for AGORA and right panel (D and F) for CarveMe. Smaller environments (essential compounds plus 50 compounds) are shown in (C) and (D) and larger environments (essential compounds plus 100 compounds) are shown in (E) and (F). As in the main paper, we found a more diverse spread of interactions in larger environments.

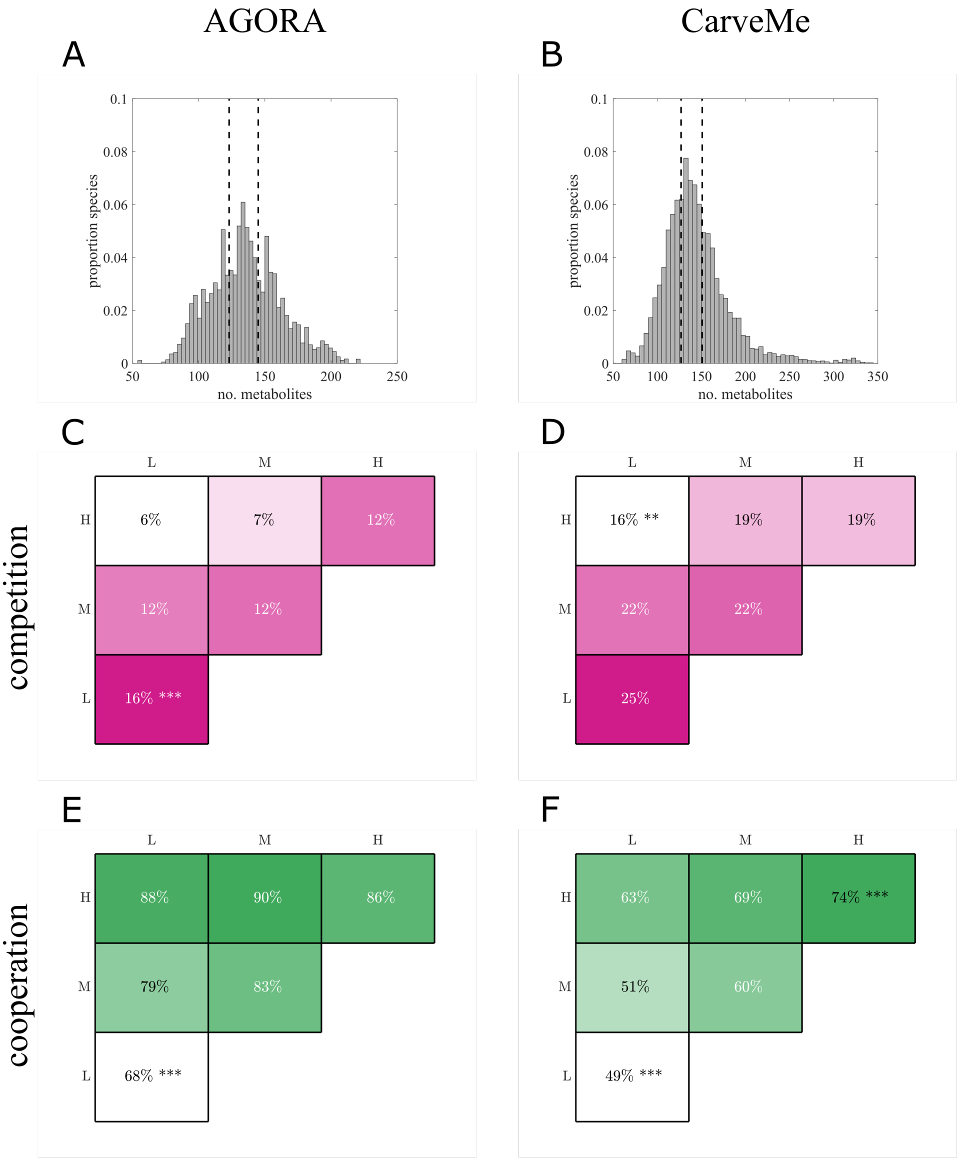


**Fig. S4. Correlation of competition or cooperation with metabolic complexity of bacteria pairs; measured by number of metabolites.** A and B) Histograms show the frequency of number of metabolites that could be used per bacteria in the AGORA (A) and CarveMe (B) collections, which can be used as a proxy for metabolic complexity. Dashed lines split the data into thirds which are used to classify 'low', 'medium', and 'high' complexity in the following plots. C–F) Box plots of the number of competitive (C and D) and cooperative (E and F) interactions per pair analysed in AGORA (C and E) and CarveMe (D and F). In all cases the classification of data by metabolic complexity with number of metabolites is statistically significantly different than if the classification was done randomly from a uniform distribution (*p-value* $< 0.001$). Stars are used to indicate whether the complexity classification with the highest and lowest observations of competitive/cooperative environments were significantly different from all other classifications using an ANOVA multiple comparison test ($***\, p < 0.001$). When both species have a low metabolic complexity, we found the highest number of competitive and fewest cooperative environments. The fewest competitive environments were found between low–high pairs, and the most cooperative environments were found for pairs with at least one high complexity species.

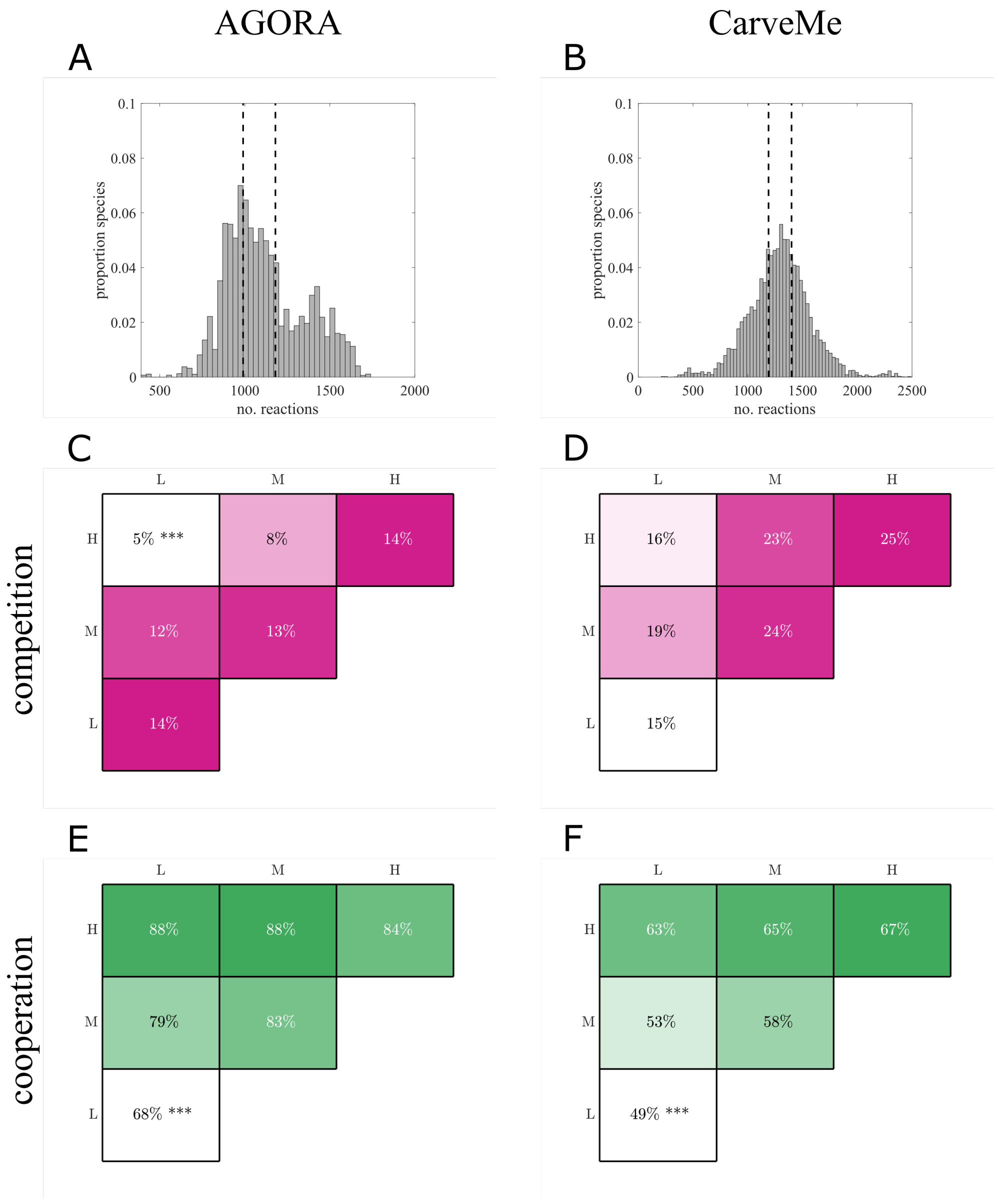


**Fig. S5. Correlation of competition or cooperation with metabolic complexity of bacteria pairs; measured by number of reactions.** A and B) Histograms show the frequency of number of reactions per bacteria in the AGORA (A) and CarveMe (B) collections, which can be used as a proxy for metabolic complexity. Dashed lines split the data into thirds which are used to classify ‘low’, ‘medium’, and ‘high’ complexity in the following plots. C–F) Box plots of the number of competitive (C and D) and cooperative (E and F) interactions per pair analysed in AGORA (C and E) and CarveMe (D and F). In all cases the classification of data by metabolic complexity with number of reactions is statistically significantly different than if the classification was done randomly from a uniform distribution (*p-value* $< 0.001$). Stars are used to indicate whether the complexity classification with the highest and lowest observations of competitive/cooperative environments were significantly different from all other classifications using an ANOVA multiple comparison test ($*** \ p < 0.001$ and $** \ p < 0.01$). The trends for cooperative environments with metabolic complexity here qualitatively agree with Figs. S4 E and F where complexity was measured by number of metabolites. However, the highest number of competitive environments was now found between high–high pairs and the pairings with fewest competitive environments differed between the two collections.

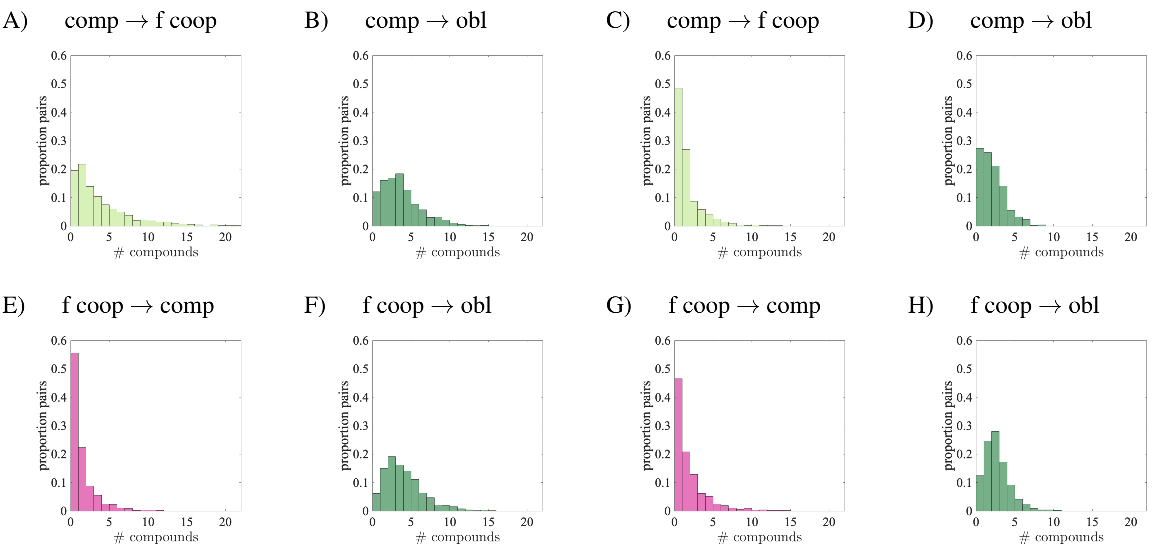


**Fig. S6. Frequency single compounds cause interaction switches.** Histograms show how often the removal of a single compound can cause different interaction switches (from competition to facultative cooperation (a and c), or obligate interactions (b and d); facultative cooperation to competition (e and g), or obligate interactions (f and h)), corresponding to the plots in Fig. 4. Left panels (a, b, e, and f) show results for AGORA and right panels (c, d, g, h) for CarveMe; the two collections appear qualitatively similar for most switches except competition to facultative cooperation where there are more compounds that can cause such switches per AGORA pairs. In all cases, except facultative cooperation to competition in AGORA, over half of the pairs considered have at least one compound that can cause the given interaction switch.

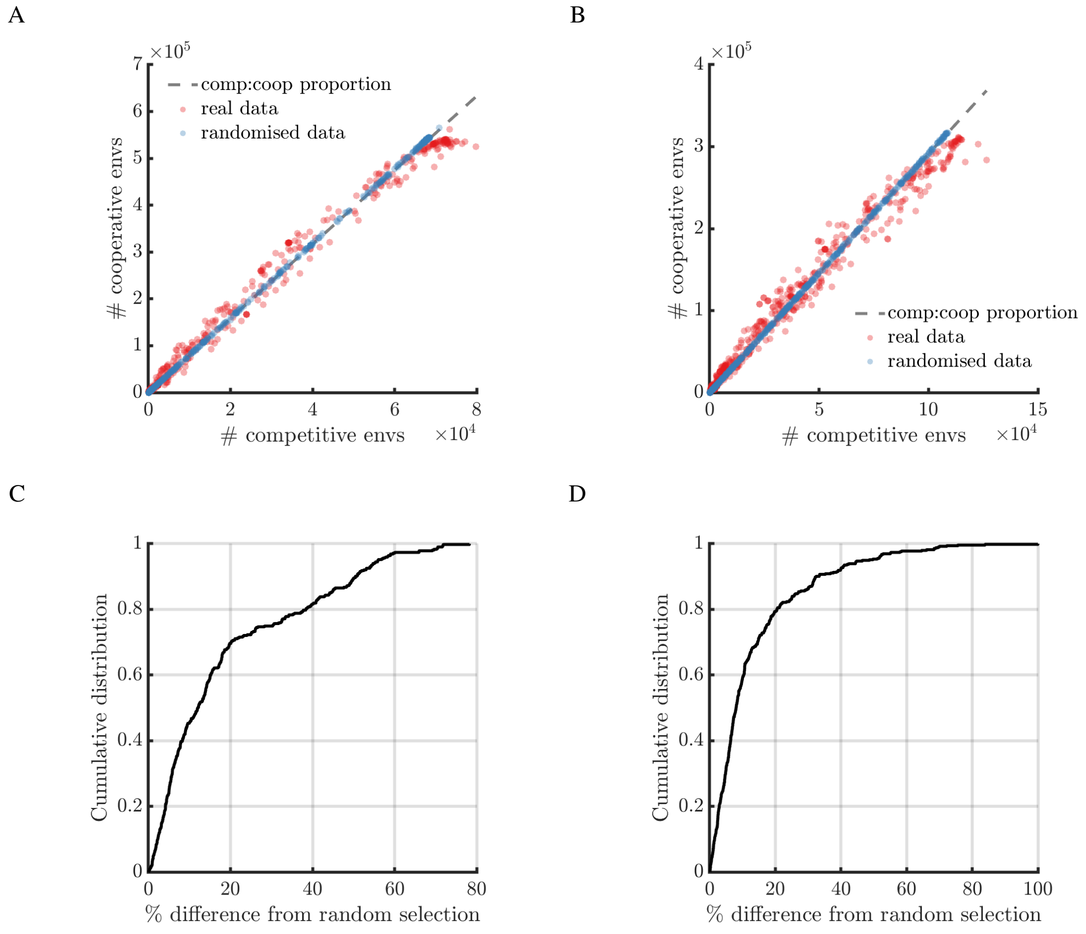


**Fig. S7. Compound appearance in competitive *vs* cooperative environments.** A and B) Plots show the number of appearances of each compound in competitive vs cooperative environments in AGORA (A) and CarveMe (B). Grey dashed line indicates the proportion of competitive to cooperative environments found (11.2% in AGORA and 25.6% in CarveMe); red dots show the real data from the environments found (939656 environments in AGORA and 804379 in CarveMe); blue dots show simulation data if the identified environments are randomly assigned competitive or cooperative with a probability given by the proportion of such environments found overall. The majority of compounds (97.4% AGORA, 95.8% CarveMe) appeared in a higher proportion of competitive or cooperative environments than by independent random sampling (*p-value* $< 0.001$). C and D) Plots show the cumulative distribution function of compound appearance biases in AGORA (C) and CarveMe (D); 68.2% compounds have a less that 20% bias away from equal probability of competitive or cooperative environments in AGORA and 78.4% in CarveMe. These show that the median bias is only 12.1% in AGORA and 8.0% in CarveMe.

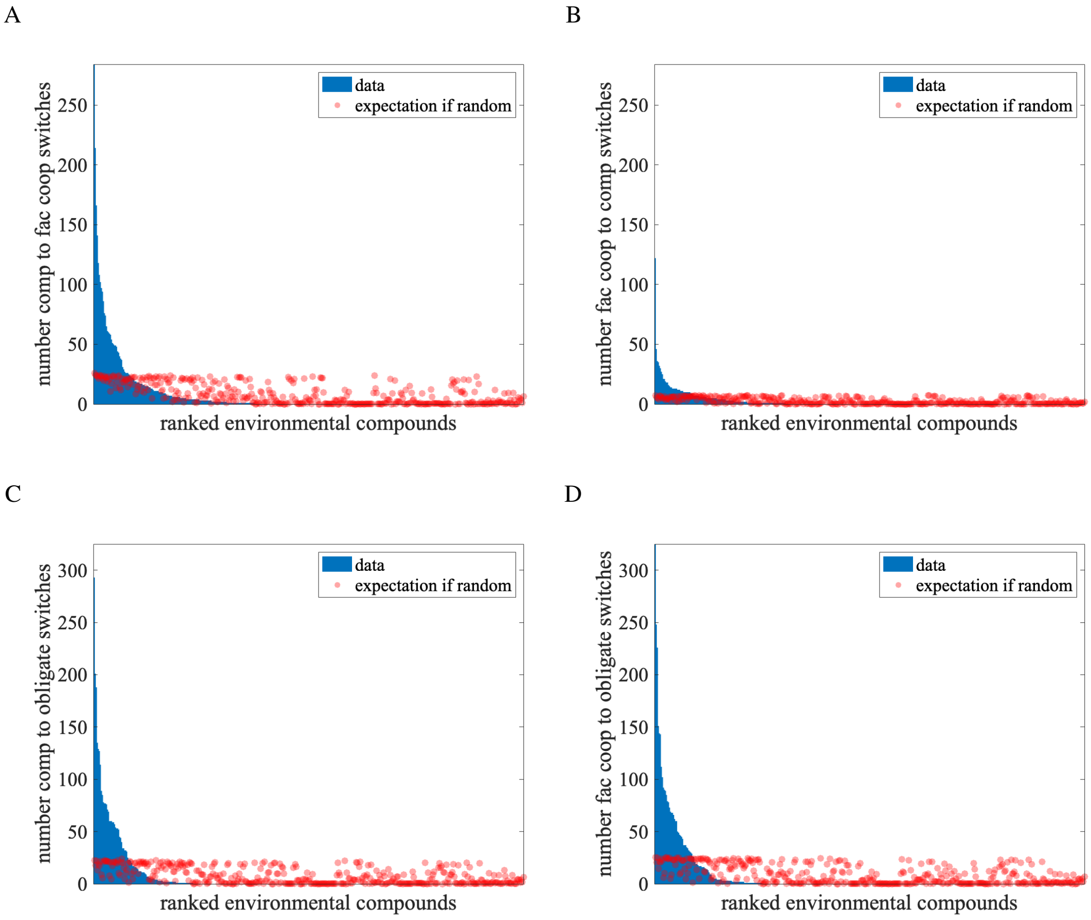


**Fig. S8. Compounds that cause switches in interactions between 1000 pairs in AGORA.** Blue bars show the data for compounds from environments generated by our search algorithm. Red dots show the expected number of switches that would be caused if it was by random chance; variation arises from the different number of environments that compounds are present in. A) Switches from competitive to facultative cooperative environments are caused by 38% of compounds. B) Switches from facultative cooperative to competitive environments are caused by 30% of compounds. C) Switches from competitive to obligate environments are caused by 23% of compounds. D) Switches from facultative cooperative to obligate environments are caused by 25% of compounds. Across these figures, only 45% compounds are responsible for all the switches observed between competitive and cooperative environments.

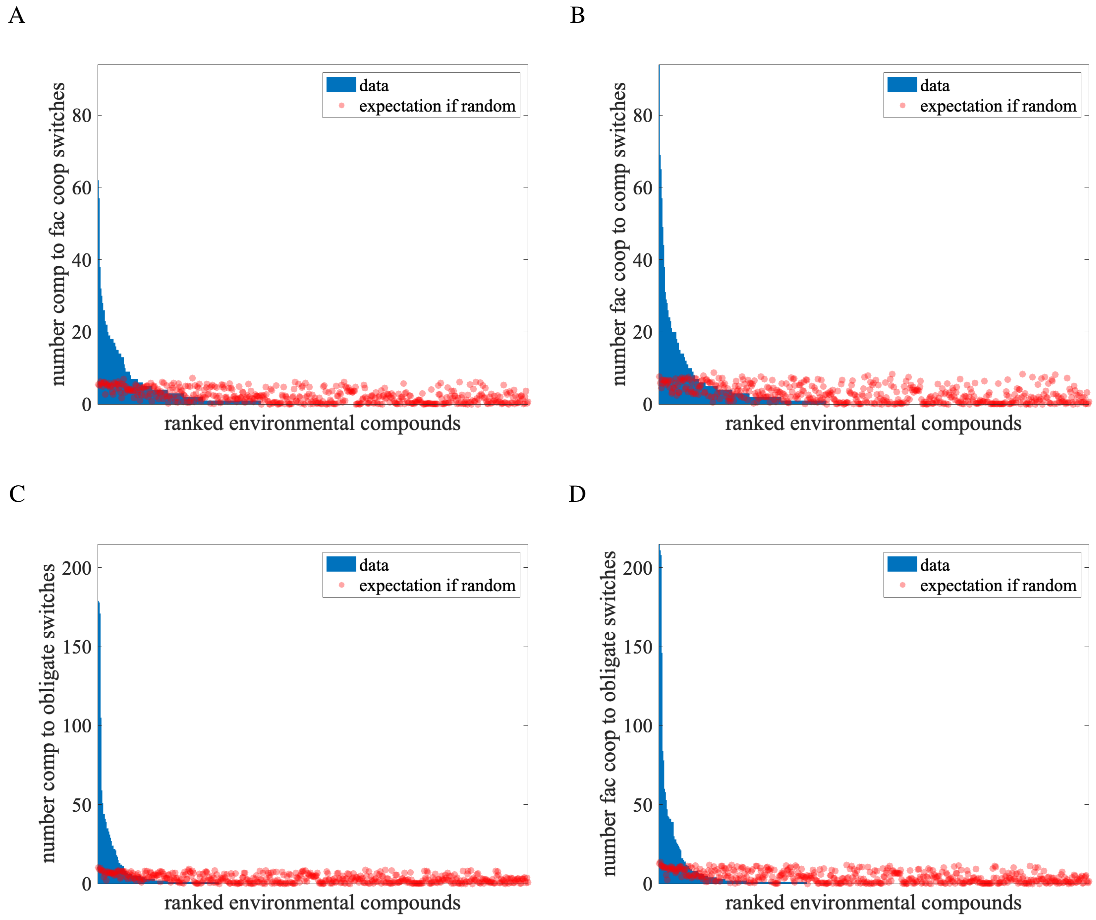


**Fig. S9. Compounds that cause switches in interactions between 1000 pairs in CarveMe.** Blue bars show the data for compounds from environments generated by our search algorithm. Red dots show the expected number of switches that would be caused if it was by random chance; variation arises from the different number of environments that compounds are present in. A) Switches from competitive to facultative cooperative environments are caused by 38% of compounds. B) Switches from facultative cooperative to competitive environments are caused by 39% of compounds. C) Switches from competitive to obligate environments are caused by 28% of compounds. D) Switches from facultative cooperative to obligate environments are caused by 34% of compounds. Across these figures, 58% compounds are responsible for all the switches observed between competitive and cooperative environments.

**Table S2. Compounds which cause transitions in AGORA.** Top ten compounds responsible for transitions a) competition to facultative cooperation; b) facultative cooperation to competition; c) competition to obligate; d) facultative cooperation to obligate in AGORA pairs considered in Fig. S8.

**(a)** Competition to facultative cooperation

| Compound | % switches |
|---|---|
| L-threonine | 8% |
| proton | 6% |
| water | 5% |
| glycylleucine | 4% |
| oxygen | 4% |
| L-glutamate(1-) | 3% |
| L-aspartate(1-) | 3% |
| L-asparagine | 3% |
| acetaldehyde | 3% |
| L-argininium(1+) | 3% |

**(b)** Facultative cooperation to competition

| Compound | % switches |
|---|---|
| oxygen | 12% |
| cytidine | 5% |
| nitrate | 4% |
| 2-oxobutanoate | 4% |
| glycylleucine | 3% |
| L-threonine | 3% |
| L-argininium(1+) | 3% |
| Adenosine | 3% |
| nitrite | 2% |
| L-alanyl-L-threonine | 2% |

**(c)** Competition to obligate

| Compound | % switches |
|---|---|
| nicotinate | 10% |
| hydrogen phosphate | 7% |
| L-valine | 6% |
| L-tryptophan | 4% |
| L-isoleucine | 4% |
| L-argininium(1+) | 4% |
| L-asparagine | 4% |
| octadecanoate | 3% |
| riboflavin | 3% |
| L-histidine | 3% |

**(d)** Facultative cooperation to obligate

| Compound | % switches |
|---|---|
| nicotinate | 9% |
| hydrogen phosphate | 7% |
| L-valine | 6% |
| L-tryptophan | 4% |
| L-isoleucine | 4% |
| L-argininium(1+) | 4% |
| L-asparagine | 3% |
| L-lysinium(1+) | 3% |
| riboflavin | 3% |
| octadecanoate | 3% |

**Table S3. Compounds which cause transitions in CarveMe.** Top ten compounds responsible for transitions a) competition to facultative cooperation; b) facultative cooperation to competition; c) competition to obligate; d) facultative cooperation to obligate in CarveMe pairs considered in Fig. S9.

**(a)** Competition to facultative cooperation

| Compound | % switches |
|---|---|
| L-arginine | 5% |
| fumarate | 5% |
| water | 3% |
| hydrogen sulfide | 3% |
| nitrite | 3% |
| glycerol 3-phosphate | 2% |
| L-glutamate | 2% |
| acetaldehyde | 2% |
| L-tartrate | 2% |
| nitrate | 2% |

**(b)** Facultative cooperation to competition

| Compound | % switches |
|---|---|
| nitrate | 7% |
| 4-aminobutanoate | 5% |
| 2-oxoglutarate | 5% |
| L-glutamate | 4% |
| D-serine | 4% |
| putrescine | 3% |
| nitrite | 3% |
| reduced glutathione | 2% |
| L-glutamine | 2% |
| nitric oxide | 2% |

**(c)** Competition to obligate

| Compound | % switches |
|---|---|
| copper 2+ | 11% |
| iron 3+ | 11% |
| iron 2+ | 11% |
| benzoate | 7% |
| L-arginine | 4% |
| hydrogen phosphate | 3% |
| L-asparagine | 3% |
| 2-phosphoglycolate | 3% |
| salmochelin-S4-Fe-III | 3% |
| oxygen | 2% |

**(d)** Facultative cooperation to obligate

| Compound | % switches |
|---|---|
| copper 2+ | 10% |
| iron 2+ | 10% |
| iron 3+ | 10% |
| benzoate | 7% |
| hydrogen phosphate | 4% |
| L-arginine | 4% |
| 2-phosphoglycolate | 3% |
| L-lysine | 3% |
| L-asparagine | 2% |
| oxygen | 2% |

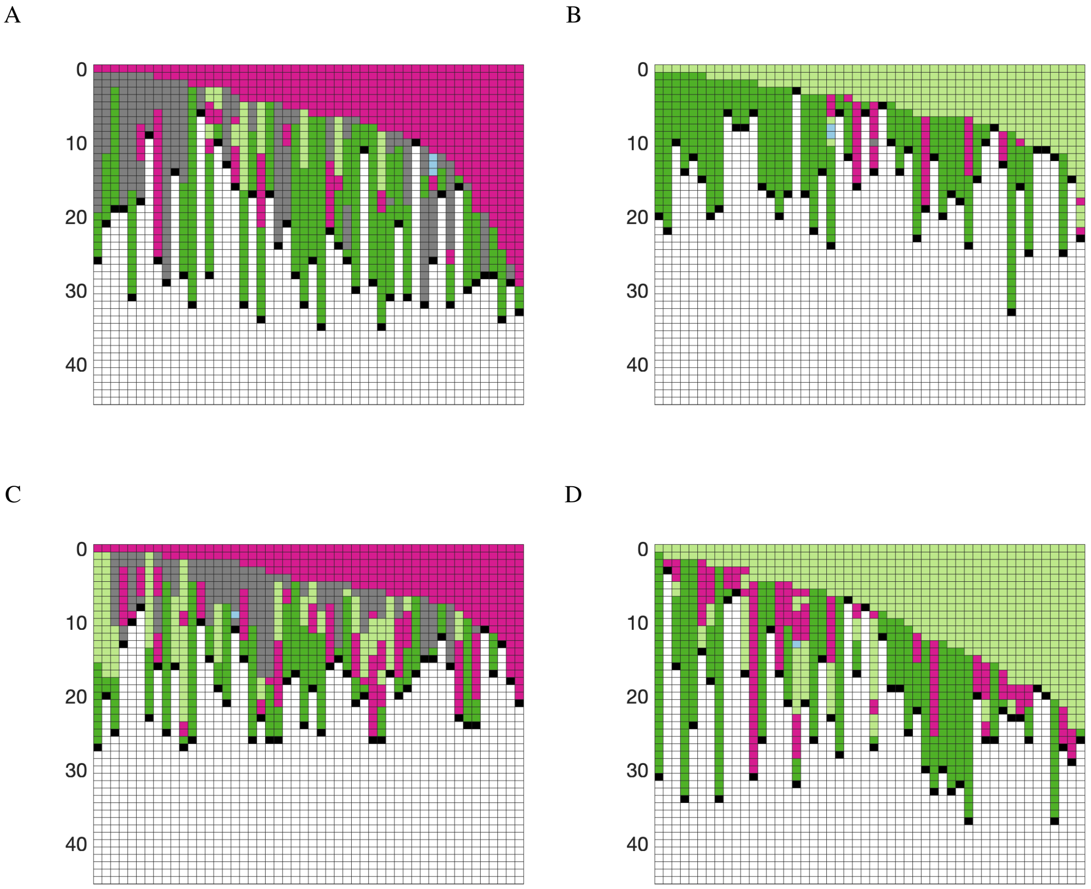


**Fig. S10. Interactions between common aquatic bacteria as compounds are removed from the environment.** Columns in the heatmaps show the interactions between pairs of bacteria as compounds are removed; going down the rows subsequent compounds are removed from the environment. Each column is a different ordering of compound removal; columns have been sorted by length of time to the first change in interaction. Colours correspond to different interactions: competition in pink; facultative cooperation in light green; one-way obligate interactions in green; two-way obligate interactions in dark green; neutral interactions in blue; no growth in black and other interactions in grey ( $(+/=)$ or $(x/=)$ ). In the left panels pairs begin in an environment for competition and in right panels for facultative cooperation. A) and B) *Prochlorococcus_marinus_str_MIT_9312* and *Pseudomonas_aeruginosa_PAO1*, C) and D) *Prochlorococcus_marinus_str_MIT_9312* and *Alteromonas_macleodii_ATCC_27126*.

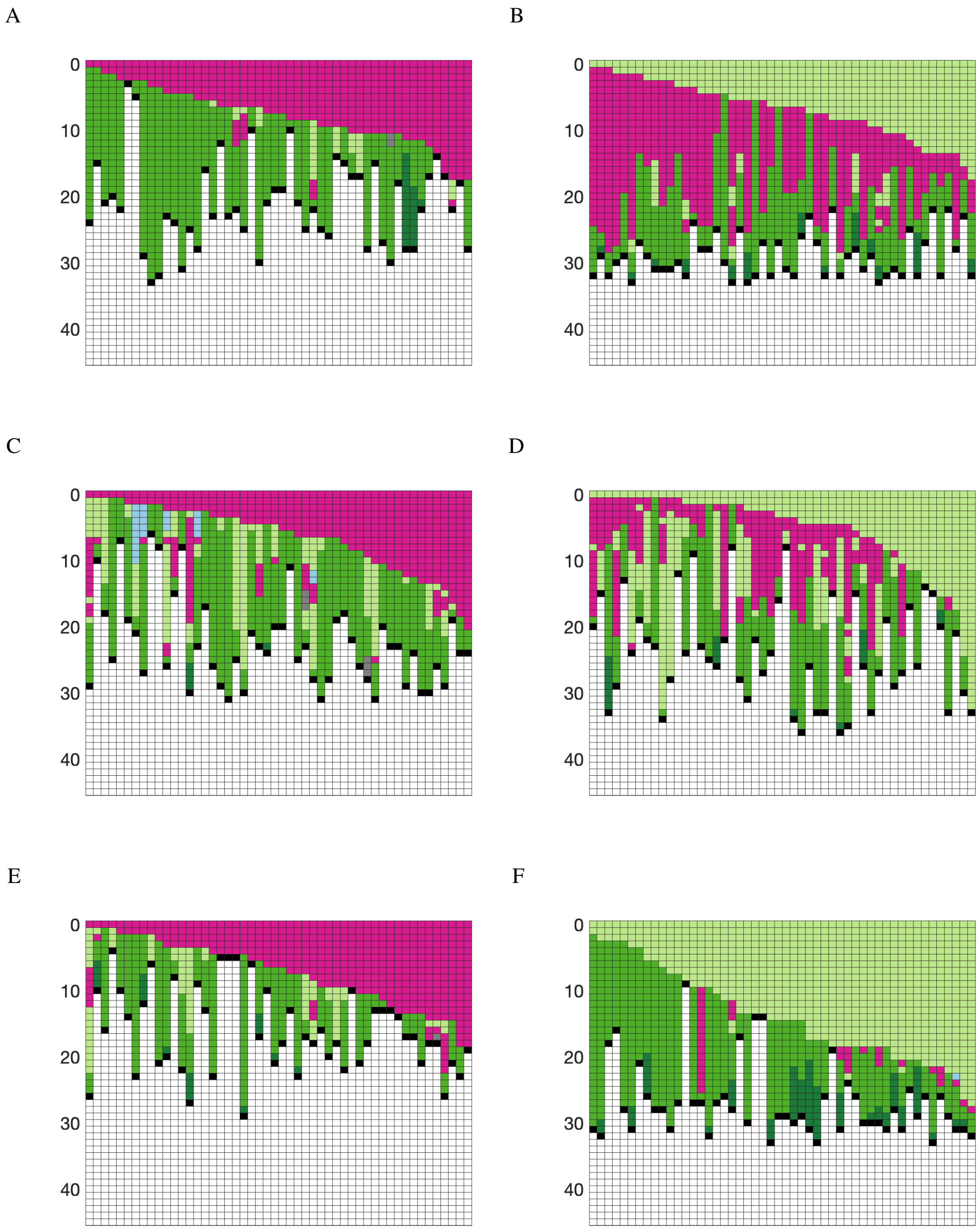


**Fig. S11. Interactions between common soil bacteria as compounds are removed from the environment.** In the left panels pairs begin in an environment for competition and in right panels for facultative cooperation. A) and B) *Bradyrhizobium_japonicum_USDA_6* and *Sphingomonas_melonis_TY*, C) and D) *Bradyrhizobium_japonicum_USDA_6* and *Methylocystis_parvus_OBBP*, E) and F) *Bradyrhizobium_japonicum_USDA_6* and *Actinoplanes_friuliensis_DSM_7358*.

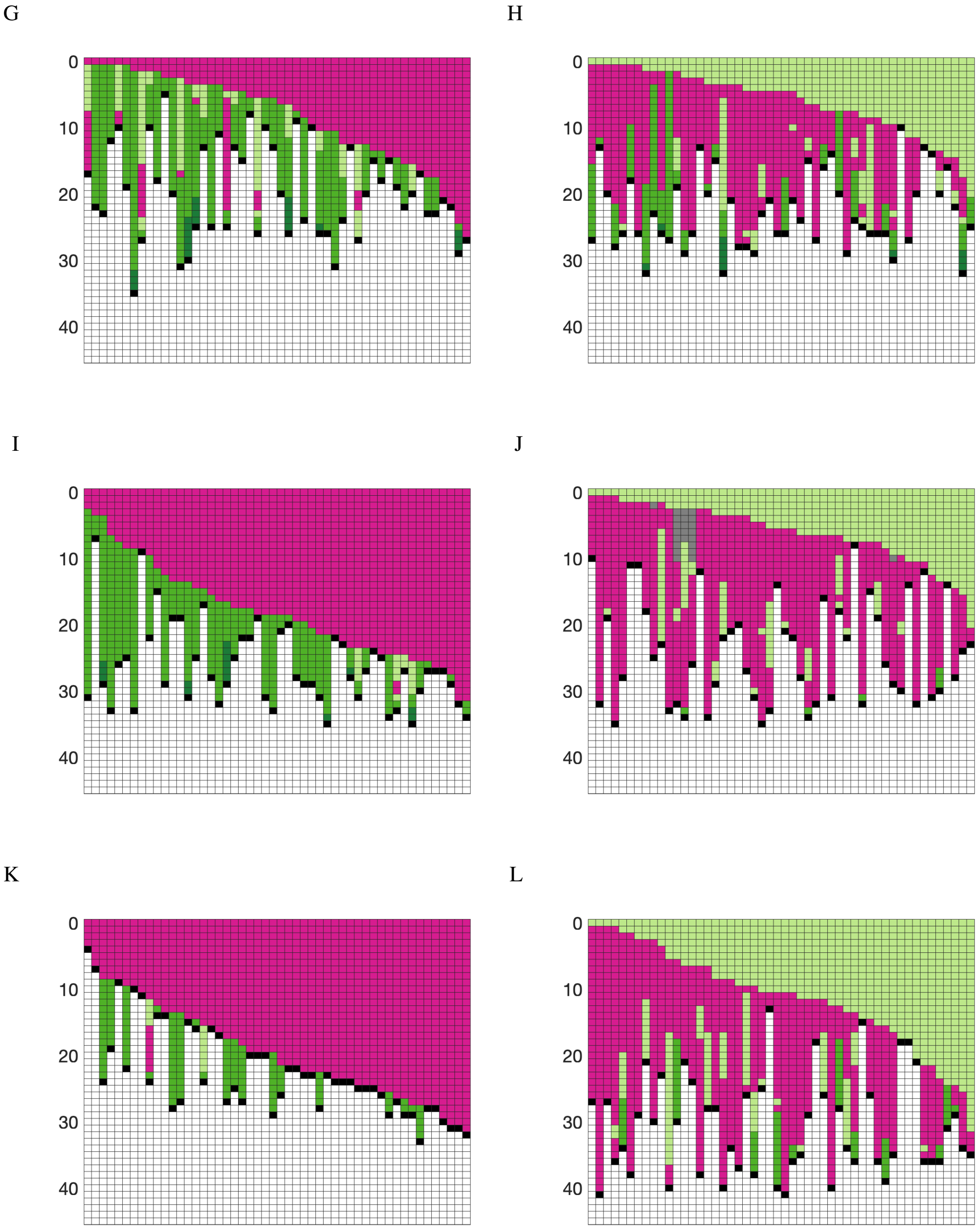


**Fig. S11.** G) and H) *Bradyrhizobium_japonicum_USDA_6* and *Blastococcus_endophyticus_DSM_45413*, I) and J) *Bradyrhizobium_japonicum_USDA_6* and *Azospirillum_brasilense_Sp_7*, K) and L) *Bradyrhizobium_japonicum_USDA_6* and *Rubrobacter_xylanophilus_DSM_9941*.

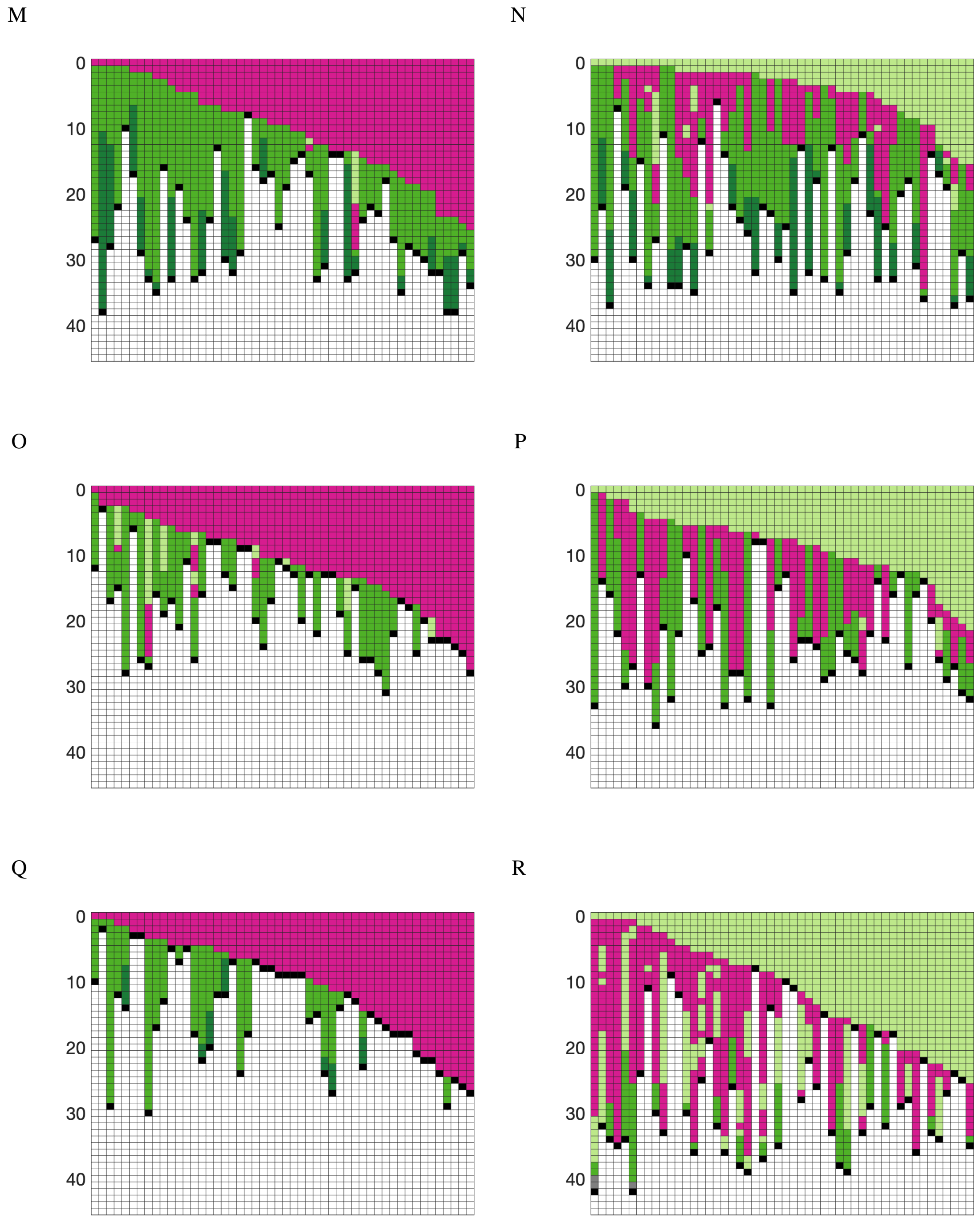


**Fig. S11.** M) and N) *Bradyrhizobium_japonicum_USDA_6* and *Mycobacterium_rhodesiae_JS60*, O) and P) *Bradyrhizobium_japonicum_USDA_6* and *Phycisphaera_mikurensis_NBRC_102666*, Q) and R) *Bradyrhizobium_japonicum_USDA_6* and *Streptomyces_coelicolor_A3_2*.

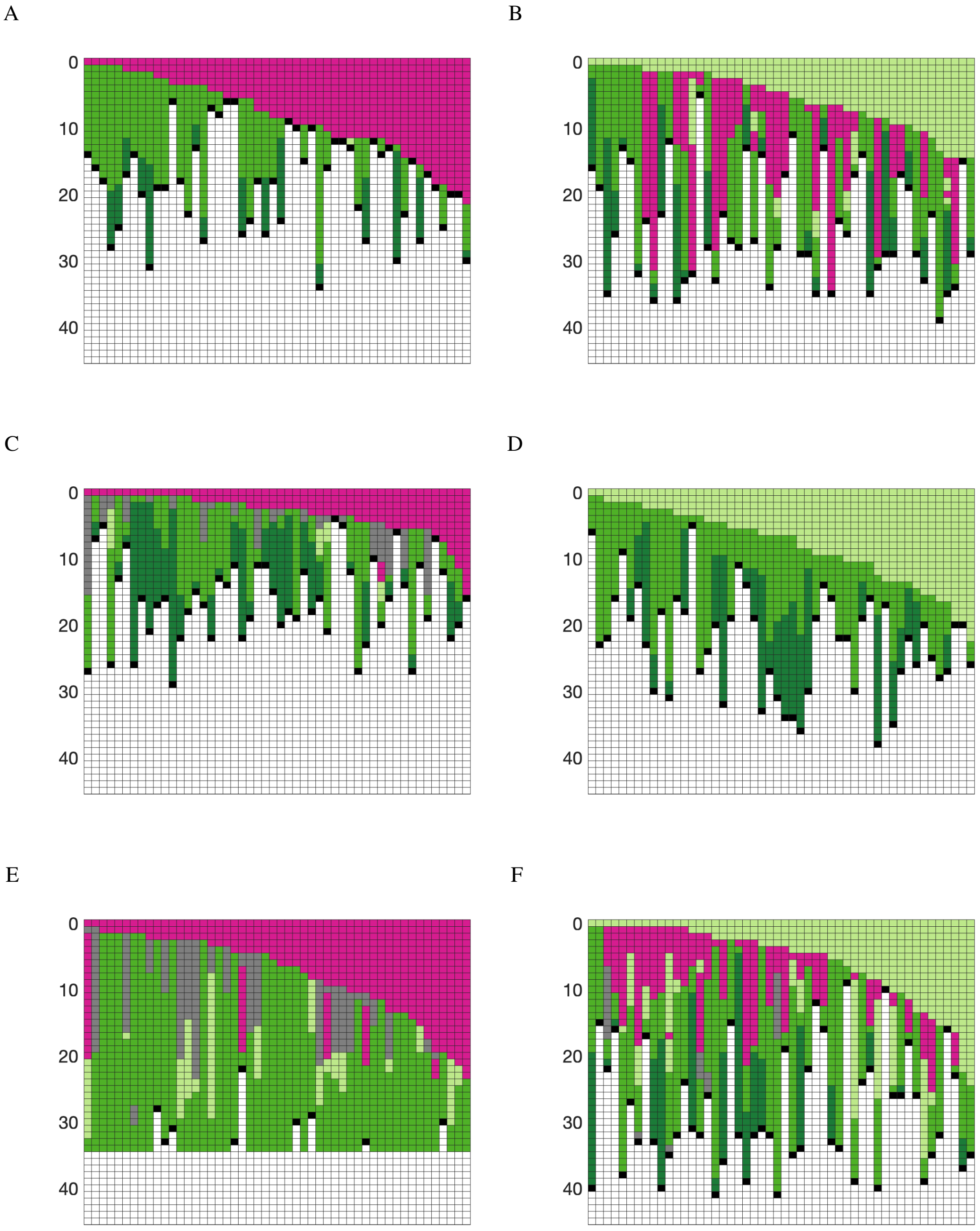


**Fig. S12. Interactions between common gut bacteria as compounds are removed from the environment (CarveMe).** In the left panels pairs begin in an environment for competition and in right panels for facultative cooperation. A) and B) *Prevotella_copri_DSM_18205* and *Bacteroides_vulgatus_ATCC_8482*, C) and D) *Prevotella_copri_DSM_18205* and *Lactobacillus_fermentum_IFO_3956*, E) and F) *Prevotella_copri_DSM_18205* and *Collinsella_aerofaciens_ATCC_25986*. We note that in some cases the set of essential compounds in an environment is sufficient for the growth of the pair, as seen in (E).

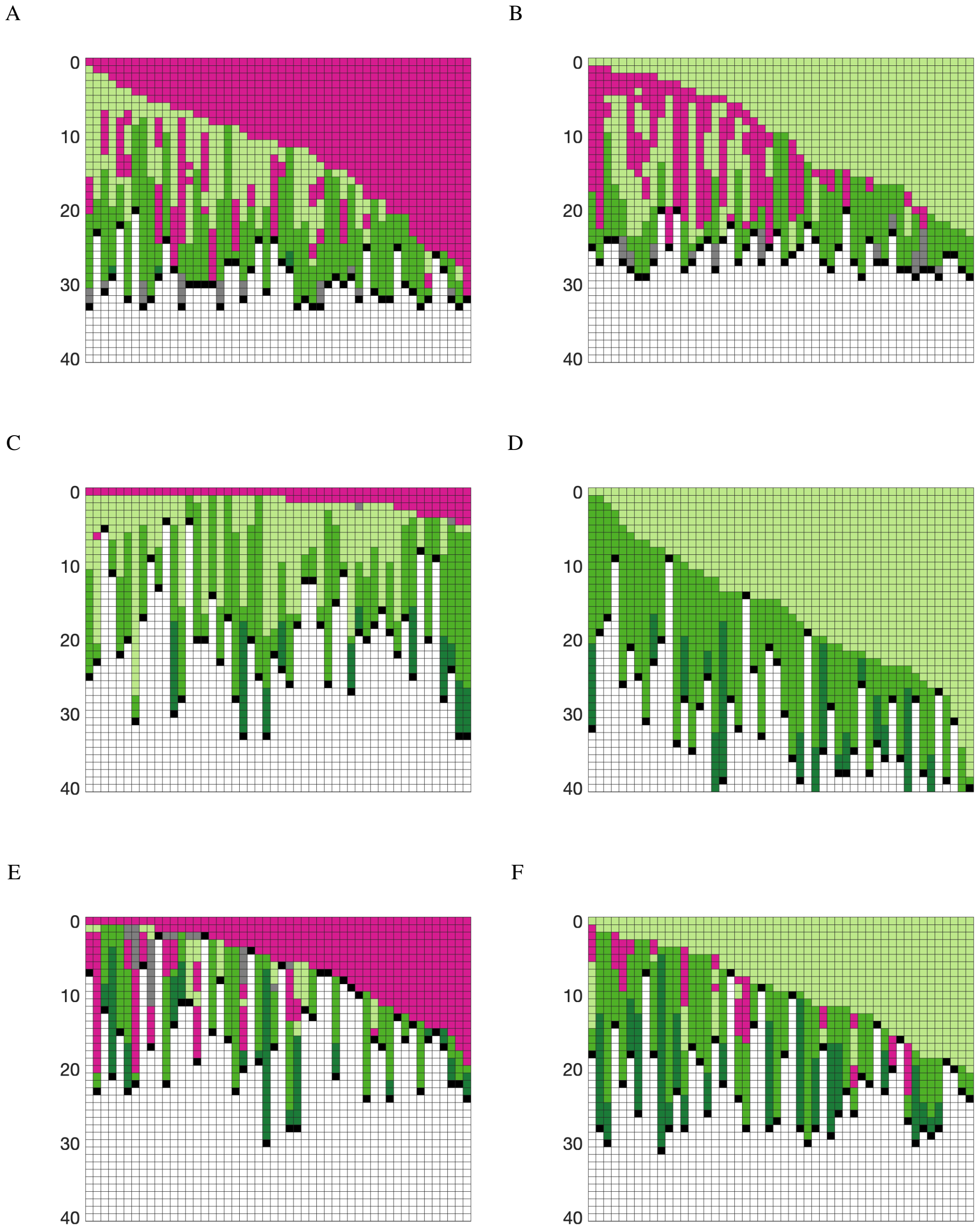


**Fig. S13. Interactions between common gut bacteria as compounds are removed from the environment (AGORA).** In the left panels pairs begin in an environment for competition and in right panels for facultative cooperation. A) and B) *Prevotella_copri_DSM_18205* and *Bacteroides_vulgatus_ATCC_8482*, C) and D) *Prevotella_copri_DSM_18205* and *Clostridium_perfringens_ATCC_13124*, E) and F) *Prevotella_copri_DSM_18205* and *Ruminococcus_gnavus_AGR2154*.

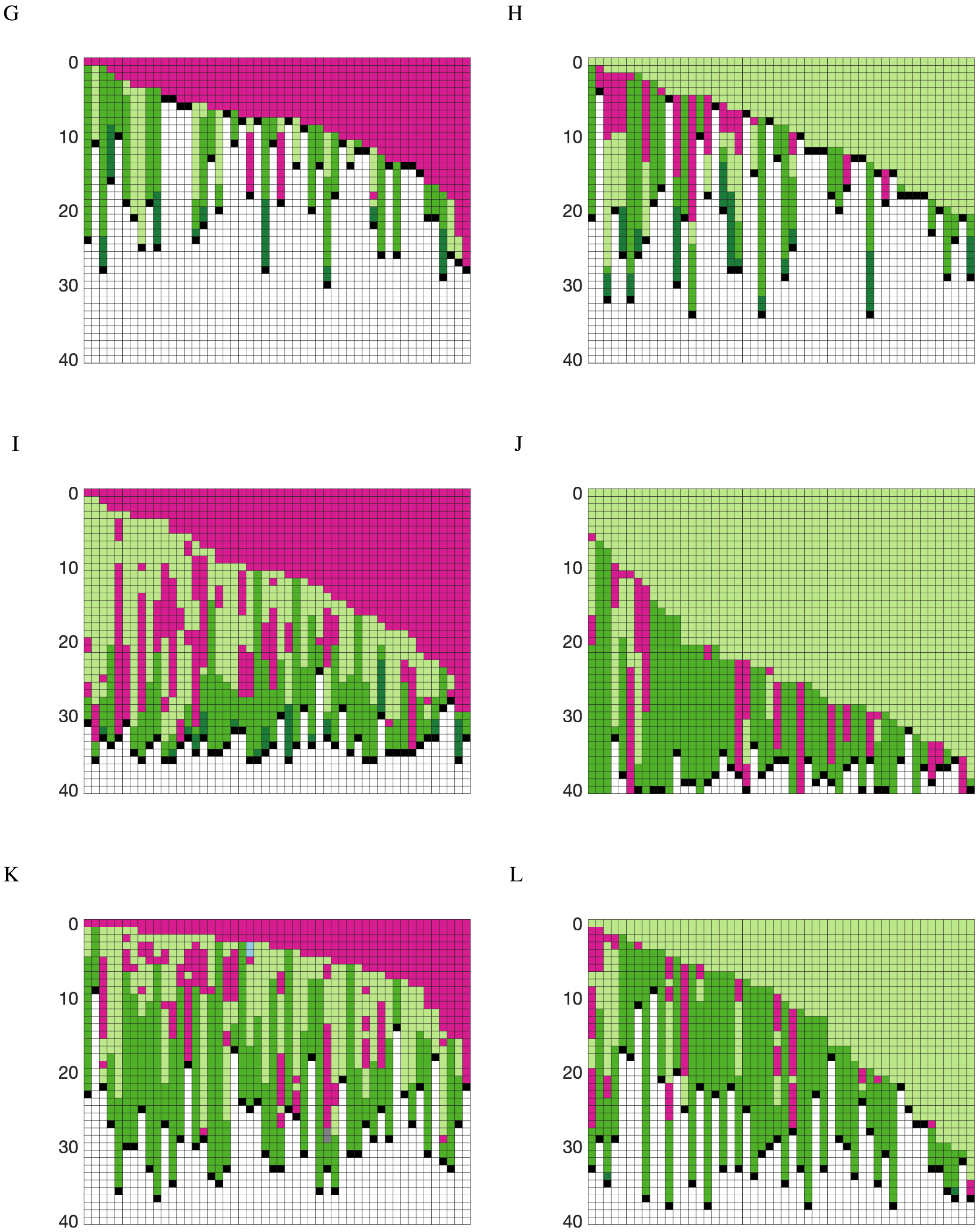


**Fig. S13.** G) and H) *Prevotella_copri_DSM_18205* and *Eubacterium_rectale_ATCC_33656*, I) and J) *Prevotella_copri_DSM_18205* and *Bifidobacterium_longum_longum_JCM_1217*, K) and L) *Prevotella_copri_DSM_18205* and *Escherichia_coli_O157_H7_str_Sakai_Sakai_substr_RIMD_0509952*.

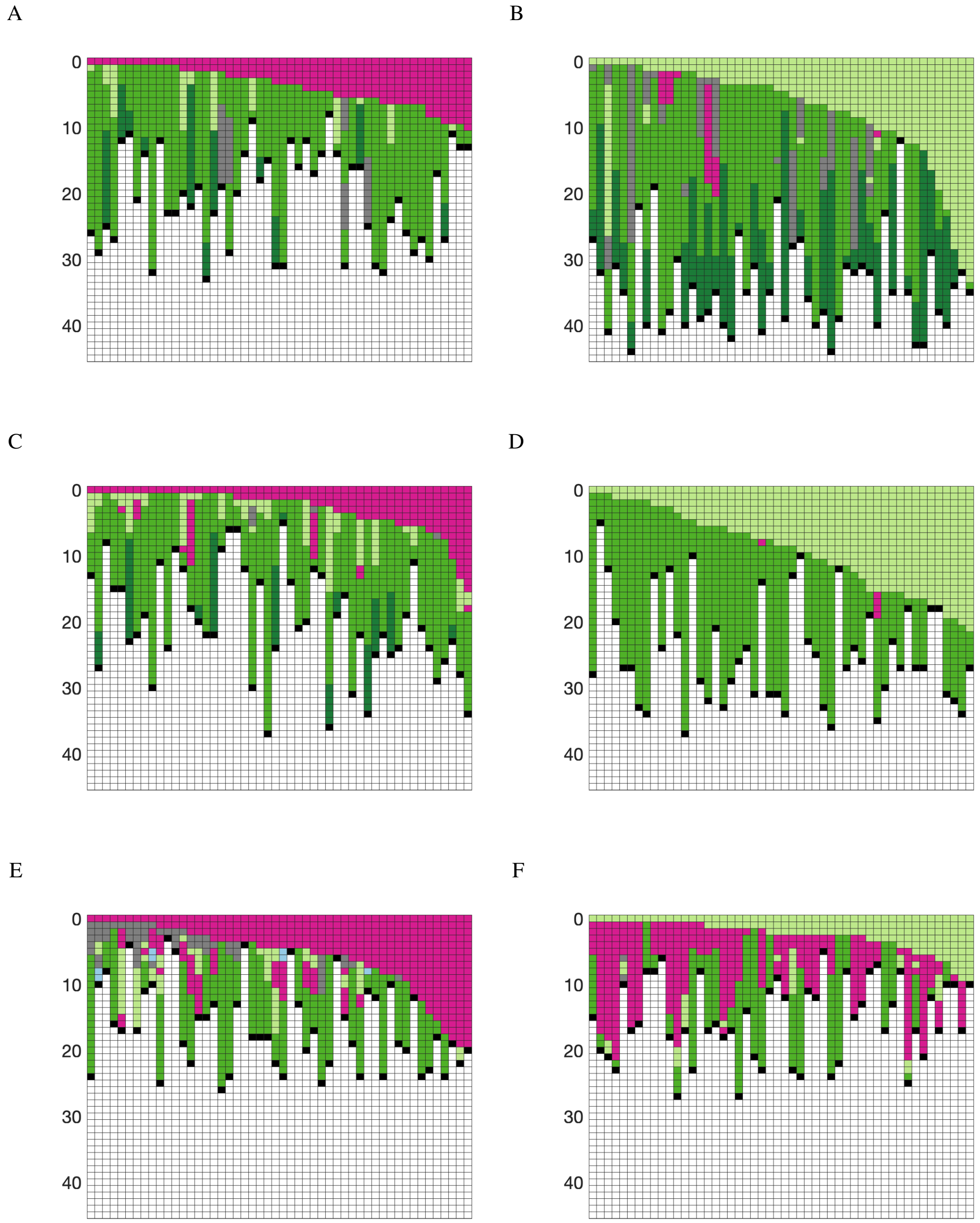


**Fig. S14. Interactions between gut bacteria *Akkermansia* and *Blautia* species as compounds are removed from the environment (CarveMe).** In the left panels pairs begin in an environment for competition and in right panels for facultative cooperation. A) and B) *Akkermansia_muciniphila_ATCC_BAA_835* and *Blautia_hansenii_DSM_20583*, C) and D) *Akkermansia_muciniphila_ATCC_BAA_835* and *Blautia_hydrogenotrophica_DSM_10507*, E) and F) *Akkermansia_muciniphila_ATCC_BAA_835* and *Blautia_obeum_2789STDY5608838*.

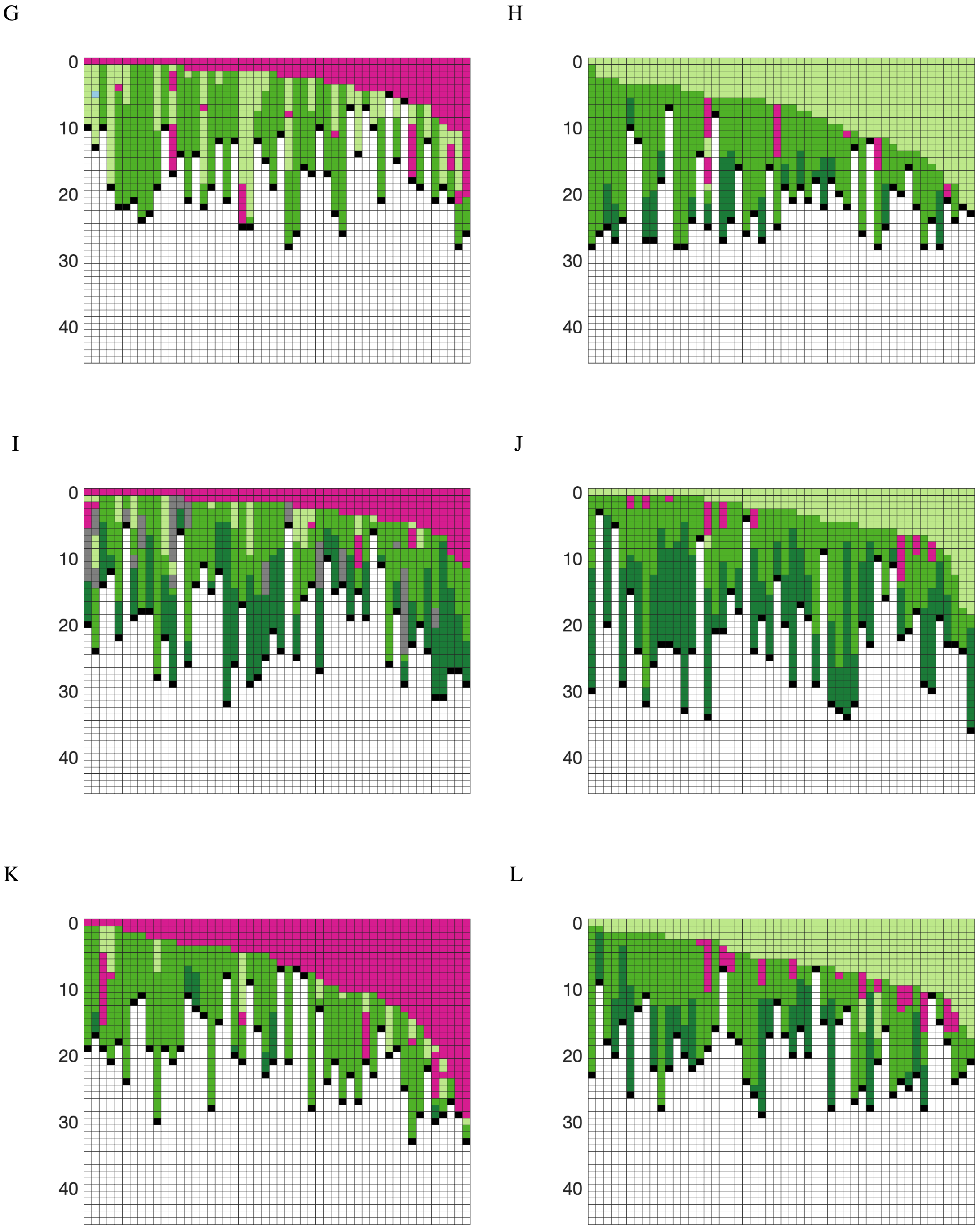


**Fig. S14.** G) and H) *Akkermansia_muciniphila_ATCC_BAA_835* and *Blautia_obeum_ATCC_29174*, I) and J) *Akkermansia_muciniphila_ATCC_BAA_835* and *Blautia_producta_ATCC_27340_DSM_2950*, K) and L) *Akkermansia_muciniphila_ATCC_BAA_835* and *Blautia_schinkii_DSM_10518*.

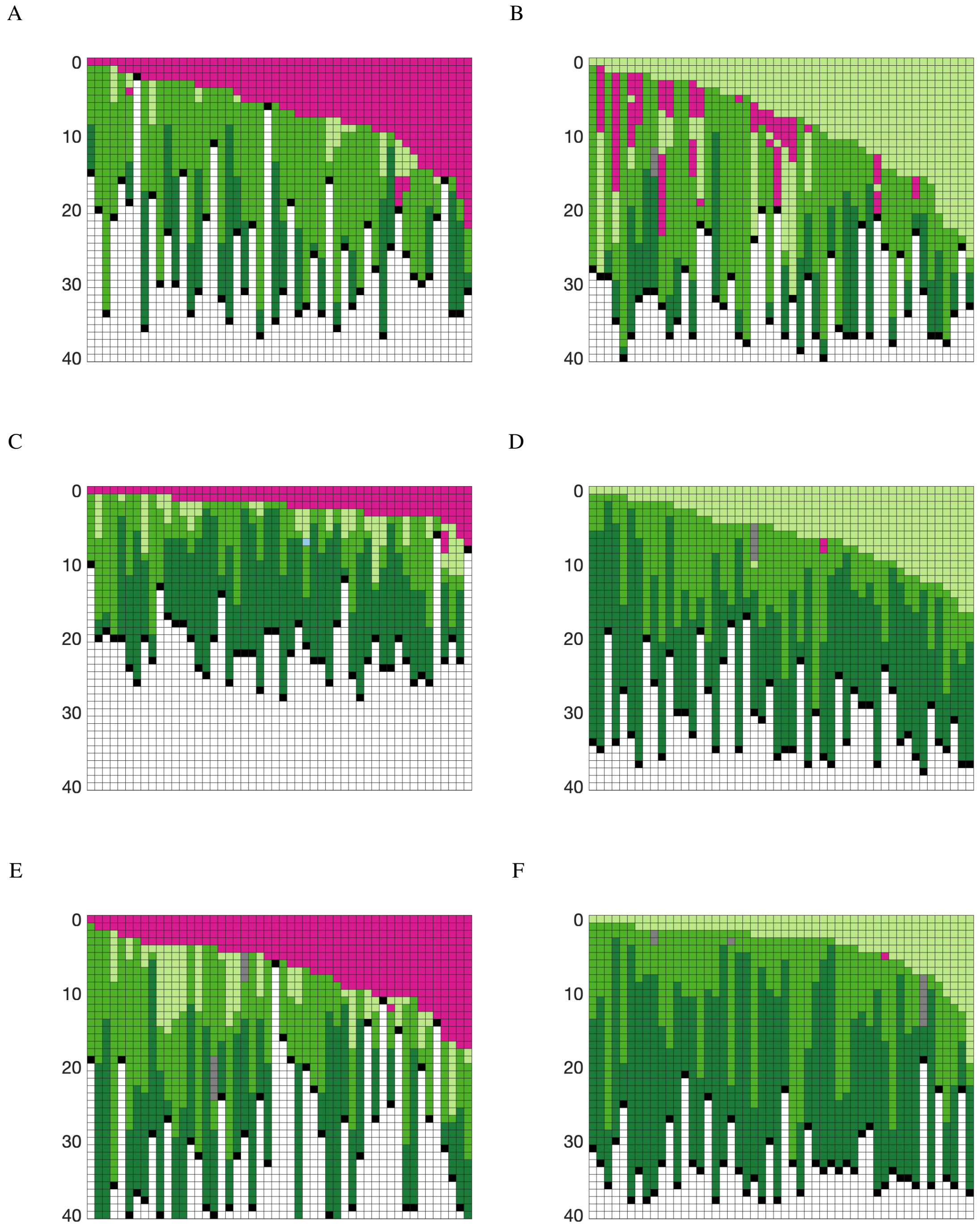


**Fig. S15. Interactions between gut bacteria *Akkermansia* and *Blautia* species as compounds are removed from the environment (CarveMe).** In the left panels pairs begin in an environment for competition and in right panels for facultative cooperation. A) and B) *Akkermansia_muciniphila_ATCC_BAA_835* and *Blautia_hansenii_VPI_C7_24_DSM_20583*, C) and D) *Akkermansia_muciniphila_ATCC_BAA_835* and *Blautia_hydrogenotrophica_DSM_10507*, E) and F) *Akkermansia_muciniphila_ATCC_BAA_835* and *Blautia_obeum_ATCC_29174*.

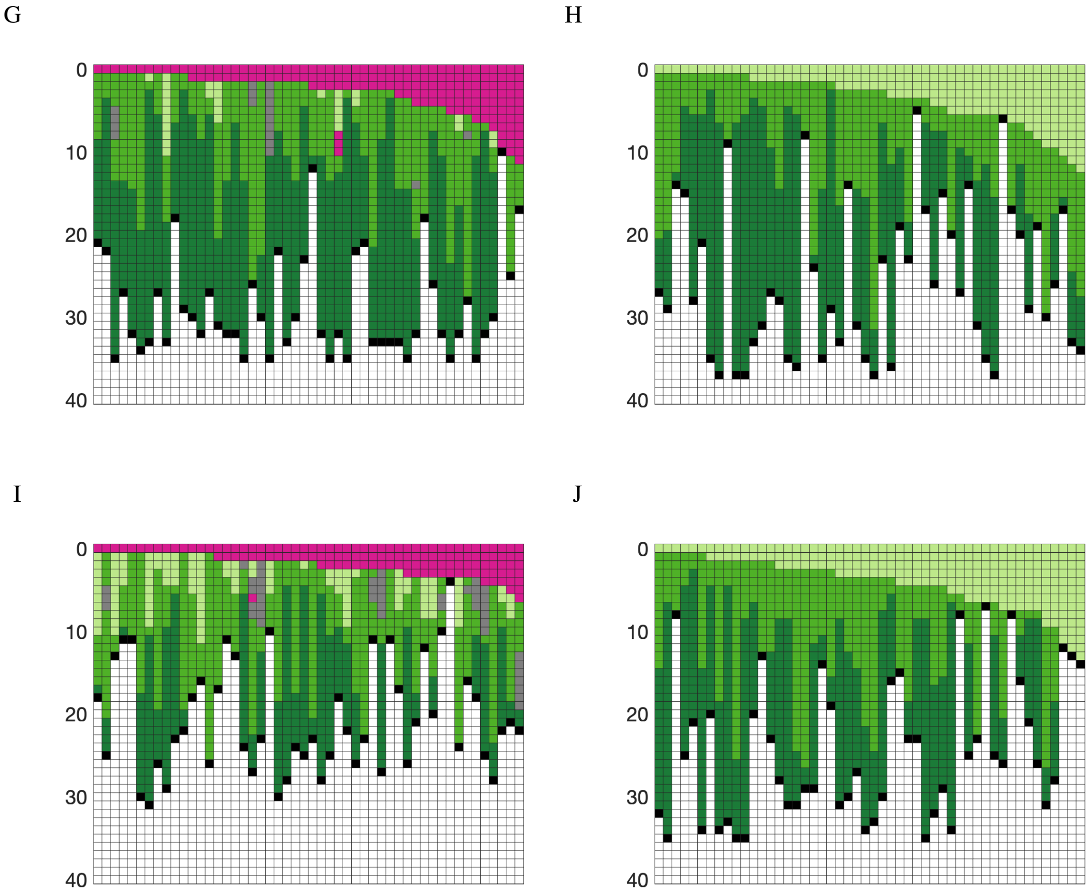


**Fig. S15.** G) and H) *Akkermansia_muciniphila_ATCC_BAA_835* and *Blautia_producta_DSM_2950*, I) and J) *Akkermansia_muciniphila_ATCC_BAA_835* and *Blautia_wexlerae_DSM_19850*.

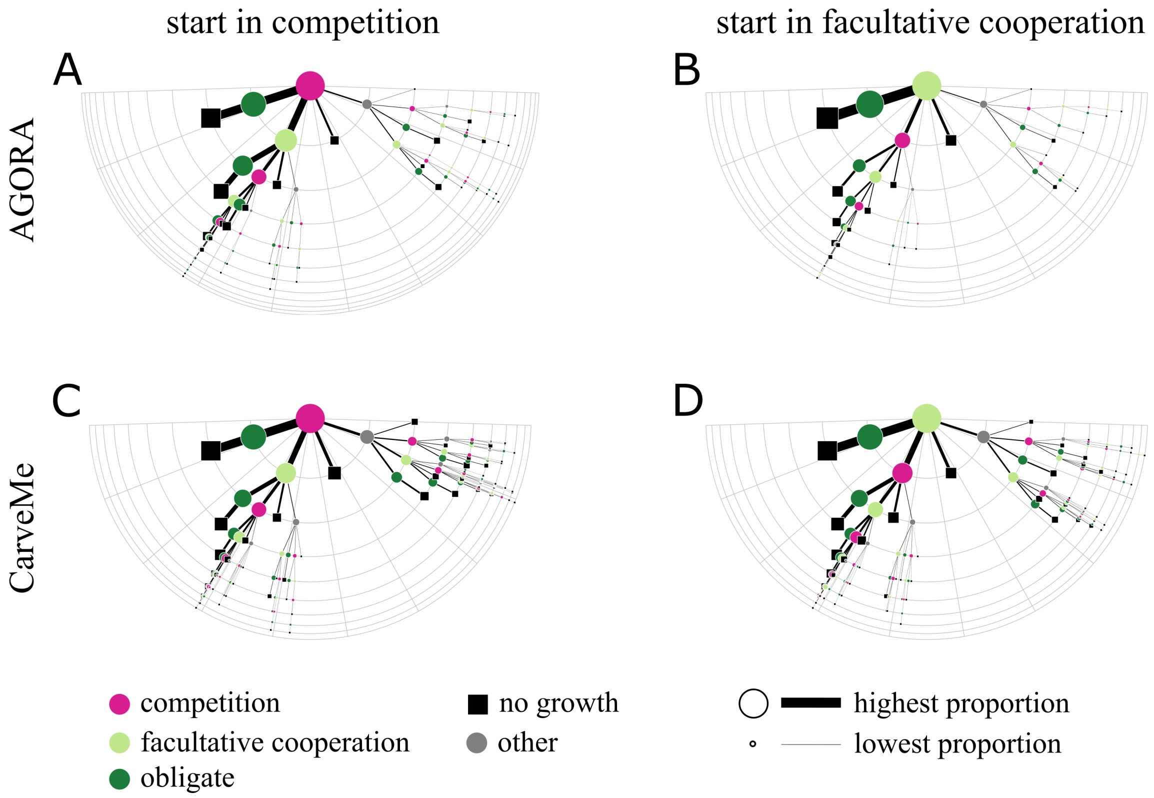


**Fig. S16. Environmental degradation summaries with all paths included.** Diagrams show the full summary of interaction changes between 300 pairs of bacteria as compounds are removed from the environment as in figure 6A–D. The top row summarises interaction paths starting in competition ((A) AGORA and (B) CarveMe), and the bottom starting in facultative cooperation (C) AGORA and (D) CarveMe). The semi-circular grid indicates the number of interaction switches. Nodes and line thickness are proportional to the square-rooted proportion of interactions taking that path; additionally paths are sorted such that the most common branch at each node is furthest to the left. All paths are included here; it was possible to observe 10 switches between interactions as the environment degraded.
